# Histopathology-assisted proteogenomics provides foundations for stratification of melanoma metastases

**DOI:** 10.1101/2023.09.29.559755

**Authors:** Magdalena Kuras, Lazaro Hiram Betancourt, Runyu Hong, Leticia Szadai, Jimmy Rodriguez, Peter Horvatovich, Indira Pla, Jonatan Eriksson, Beáta Szeitz, Bartłomiej Deszcz, Charlotte Welinder, Yutaka Sugihara, Henrik Ekedahl, Bo Baldetorp, Christian Ingvar, Lotta Lundgren, Henrik Lindberg, Henriett Oskolas, Zsolt Horvath, Melinda Rezeli, Jeovanis Gil, Roger Appelqvist, Lajos V. Kemény, Johan Malm, Aniel Sanchez, A. Marcell Szasz, Krzysztof Pawłowski, Elisabet Wieslander, David Fenyö, Istvan Balazs Nemeth, György Marko-Varga

## Abstract

Here we describe the histopathology-driven proteogenomic landscape of 142 treatment-naïve metastatic melanoma samples. We identified five proteomic subtypes that integrate the immune and stroma microenvironment components, and associate with clinical and histopathological parameters, providing foundations for an in-depth molecular classification of melanoma. Our study shows that BRAF V600 mutated melanomas display heterogeneous biology, where the presence of an oncogene-induced senescence-like phenotype improves patient survival. Therefore, we propose a mortality-risk-based stratification, which may contribute to a more personalized approach to patient treatment. We also found a strong association between tumor microenvironment composition, disease progression, and patient outcome supported by single-cell omic signatures that point to straightforward histopathological connective tissue-to-tumor ratio assessment for better informed medical decisions. A melanoma-associated signature of single amino acid variants (SAAV) responsible for remodeling the extracellular matrix was uncovered together with SAAV-derived neoantigen candidates as targets of anti-tumor immune responses. Overall, this study offers comprehensive stratifications of melanoma metastases that may help develop tailored strategies for diagnosing and treating the disease.

## INTRODUCTION

Malignant melanoma is the most aggressive skin cancer with high metastatic potential ^1,2^ and is responsible for 80% of skin cancer-related deaths ^3^. Due to its heterogeneous nature and unpredictable metastatic progression, melanoma poses a significant challenge to the healthcare system ^4^.

In the past decades, genomic studies revealed the importance of the oncogenic driver mutations in BRAF, NRAS, NF1, and KIT genes in tumor development ^3,5–8^. The oncogenic mechanisms of the PTEN/phosphoinositol-3-kinase signaling pathway, RAC1, CDKN2A, telomerase reverse transcriptase promoter mutations, and several molecular alterations specific to mucosal and chronically sun-damaged melanomas have also been implicated in progression ^9^.

Recently, multiple effective treatment options have been developed for metastatic melanoma by targeting the tumor cells using kinase inhibitor therapies, or by targeting the surrounding tumor microenvironment (TME) by the indirect action of immune checkpoint blockade-based therapies. However, many patients do not respond to immunotherapy, and patients on targeted therapy eventually progress ^10,11^.

Mutation- and transcriptomic-based melanoma classifications have been proposed to improve clinical management ^7,12–17^. However, proteomic-based approaches with detailed clinical data are lacking. Here we studied a cohort of treatment-naïve lymph node metastases collected before the era of immune checkpoint and targeted therapies. This sets apart our study from others, often focusing on therapy responder-non-responder relationships. We hypothesized that a proteogenomic and histopathological analysis of these samples allows for a unique understanding of melanoma biology associated with the natural disease progression. In addition, protein markers and phenotype-genotype correlations derived from the study can be considered independent prognostic factors of therapy. The present study provides a comprehensive stratification of melanoma metastases including 1) proteomic subtypes that integrate the tumor immune and stroma components, and associate with clinical and histopathological features; 2) survival-based classifications of BRAF V600 mutated metastases; and 3) subgroups based on the surrounding TME which contribute to disease progression and clinical outcome. We also identified a landscape of melanoma-associated SAAV responsible for extracellular matrix remodeling together with SAAV-derived neoantigens as potential targets of anti-tumor immune responses. Finally, our study exposes complex traits in the spatial expression of survival protein markers in metastases and primary tumors.

## RESULTS

### Proteogenomic map and classification of treatment-naïve melanoma metastases

Global proteomic and phosphoproteomic analyses were performed on 142 metastatic melanoma samples from lymph nodes (126), cutaneous (1), subcutaneous (7), visceral (3), and uncharacterized (5) origin (Figure S1A). The analysis was supplemented with clinical data and histopathological assessment of nearby cancer tissue (Figure S1B and Table S1A). In addition, the proteomic and phosphoproteomic data were integrated with a previously published transcriptomic dataset from matched tumors ^12^. A total of 12,695 proteins and 45,356 phosphosites were quantified, with 8,124 proteins and 4,644 phosphosites quantified in every sample (STAR Methods and Table S2A-D). Several metrics were analyzed to assess the reliability of the proteomic workflow and the quality of the generated data (STAR METHODS and Figure S1B-H).

Two independent studies have classified melanoma tumors based on transcript levels ^7,13^. Since proteins are the functional entities in the cell, we investigated whether proteomic data could improve the classification of melanoma. Using consensus clustering (STAR Methods), we identified five major melanoma subtypes, which were classified as extracellular (EC, n = 23), extracellular-immune (EC-Im, n = 26), mitochondrial (Mit, n = 30), mitochondrial-immune (Mit-Im, n = 23), and extracellular-mitochondrial (EC-Mit, n = 16) according to their characteristic enrichment in Gene Ontology (GO) terms and Kyoto Encyclopedia of Genes and Genomes (KEGG) pathways (Figure 1A and Table S3).

**Figure 1.**
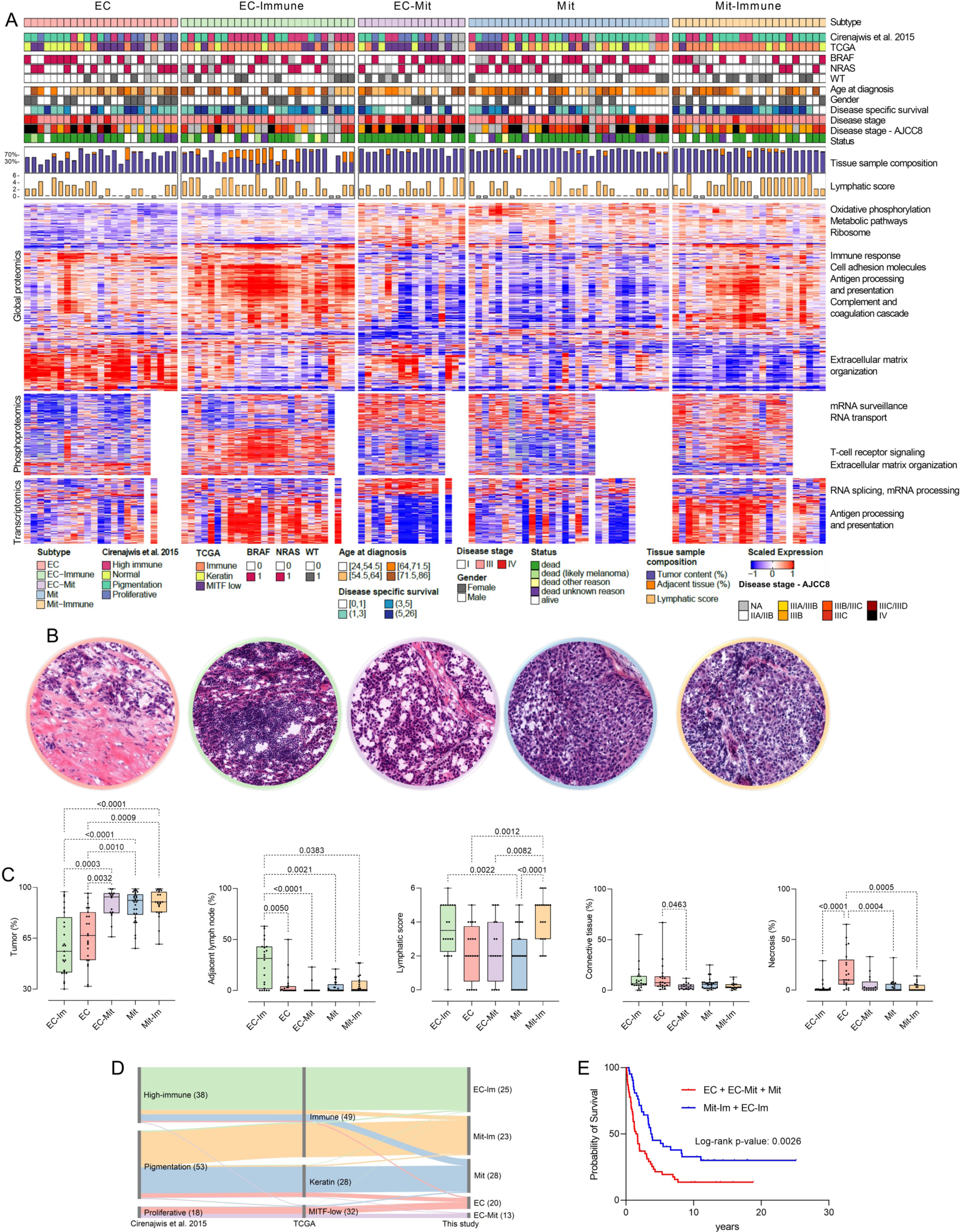
Proteomic classification of metastatic melanoma. (A) Overview of tumors grouped according to the proteomic subtypes established in this study, annotated with the transcriptomic classifications 7,12 together with clinical and histological data. Heatmaps of the most variable proteins (top 500, FDR < 0.005), phosphosites (top 1000, FDR < 0.05), and transcripts (top 500, FDR < 0.05) based on ANOVA across the five proteomic subtypes and enriched pathways annotation. (B) Representative histological images of the proteomic subtypes, EC (pink), EC-Im (green), EC-Mit (purple), Mit (blue), and Mit-Im (yellow). (C) Distribution of the annotated histological parameters; tumor content; adjacent lymph node; lymphatic score; connective tissue, and necrosis among the five proteomic subtypes (Kruskal-Wallis and Dunn’s multiple comparisons test). (D) Sankey diagram showing the association between proteomic (this study) and published transcriptomic subtypes. The EC-Im subtype was significantly associated with the high-immune (FDR = 0.021) and immune (FDR = 0.0095) groups. The Mit-Im subtype was significantly linked to the pigmentation (FDR = 0.021) and the immune (FDR = 0.0021) groups. In comparison, the EC-Mit was significantly associated with the proliferative (p-value = 0.014) and the MITF-low (FDR = 0.026) groups, and the Mit proteomic subtype was significantly linked to the keratin class (FDR = 0.040) from the TCGA classification, and the EC subtype was associated with the proliferative (p-value = 0.0244) and the MITF-low (p-value = 0.032) groups. (E) Disease-specific survival (DSS, time from surgical intervention to death or censoring) probability for patients with tumors in subtypes associated with long and short survival.

We found significant associations between the proteomic and the two transcriptomic subtypings ^12,15^, displaying a mutual validation of the three classifiers (Figure 1D). Metastases belonging to the immune transcriptomic subtypes were divided into the EC-Im and Mit-Im proteomic subtypes. In addition, the MITF-low and proliferative subtypes were separated into the EC and the EC-Mit subtypes, and tumors within the Pigmentation were mainly divided into the two Mit subtypes, while the Keratin transcriptomic subtype was mostly split among the Mit and EC subtypes.

Pathway enrichment analysis of the proteomic subtypes at protein and phosphoprotein levels showed upregulation of oxidative phosphorylation, ribosome, metabolic, and RNA-related pathways in the Mit, Mit-Im, and EC-Mit subtypes (Figure 1A). Pathways associated with extracellular matrix organization, complement, and coagulation pathways were enriched in EC, EC-Im, and EC-Mit subtypes, suggesting a more invasive phenotype ^18^. The EC-Im and Mit-Im subtypes displayed enrichment of immune signaling with upregulation of antigen processing and presentation and T cell receptor signaling pathways, including higher expression of PDL1 relative to Mit and EC-Mit (Figure S2A). At the transcript level, the EC-Mit subtype showed pathways of increased RNA activity, while in both immune subtypes, the antigen processing and presentation pathway was enriched.

To identify subtype signatures, we used Independent Component Analysis (ICA) to select the top ten most contributing proteins to the independent components (ICs) significantly correlated with the proteomic subtypes (Figure S2B). In the EC-Im subtype, we identified upregulation of the antigen receptor-mediated signaling pathway, emphasizing the immune system’s involvement in this subtype. Furthermore, positive cell-cell and cell-matrix adhesion regulation was seen, contributing to the extracellular matrix component of this subtype. Several proteins from the S100 family (S100P, S100A12, S100A8, and S1009) were identified in the EC subtype, suggesting their potential as markers for this subtype. In addition, enrichment of neutrophil degranulation was observed, which may contribute to poor prognosis and short overall survival in melanoma by upregulation of proliferation, angiogenesis, and matrix remodeling ^19^. The EC-Mit signature included a set of downregulated proteins, but no pathway enrichment was detected, not even when all the proteins (8,572) of this IC were submitted for enrichment analysis. In the protein signature associated with the Mit subtype, we found enrichment in important mitochondrial proteins such as OAT ^20^, VDAC2 ^21^, and SHMT2 ^22^. High expression of S100A1 was apparent, pointing to a unique and strong association between this melanoma marker and the Mit subtype. The top ten signature proteins in the Mit-Im subtype were associated with a downregulation of the complement and coagulation cascade rather than enrichment in a mitochondria-related signature or an upregulation of protective immune mechanisms. When upregulated in the tumor microenvironment, the complement and coagulation cascade may enhance tumor growth and increase metastasis, and it is suggested to contribute to epithelial-mesenchymal transition (EMT) ^23,24^. In addition, we could see an upregulation of these proteins in the EC subtype, again highlighting the molecular differences between these two subtypes.

### Proteomic subtypes associate with clinical and histological features

We found differences in histopathological features among the proteomic subtypes. (Figures 1B and 1C). The Mit-Im subtype was associated with higher lymphocyte density (Fisher exact test, FDR = 0.0003) than the non-immune subtypes. Both immune subtypes displayed a higher lymphatic score (ANOVA test, FDR < 0.01) than the rest of the metastases. Tumor cell content was higher in samples belonging to Mit-Im, Mit, or EC-Mit when compared to samples within the EC-Im and EC subtypes (Kruskal-Wallis test, FDR < 0.005). Adjacent lymph node and necrosis content were higher in the EC-Im and EC subtypes compared to the others (Kruskal-Wallis test, FDR < 0.05). The EC subtype, in particular, comprises 70% of the samples with the highest necrosis content (>20%). In contrast, connective tissue content was higher in the EC than in the EC-Mit subtype (Kruskal-Wallis test, FDR = 0.046).

The histological images of the Mit subtype metastases showed solid nests of highly aggressive tumor cells with broader eosinophilic cytoplasms and atypical nuclear features (Figure 1B). This phenotype reflects the picture often seen in more differentiated tumors. In the EC subtype, we observed an abundance of the intercellular matrix as desmoplastic stromal change with a pauci-cell matrix, typical characteristics of phenotype switching in melanoma. Mit-Im and EC-Im appear to be subclasses of the highly aggressive Mit and EC subtypes, respectively, where the presence of adaptive immune cells may indicate a more favorable clinical behavior, pointing towards a possible antitumor effect of adjacent lymphatic tissue ^25^. Histologically, the EC-Mit group also represents a dedifferentiated state, which may be an intermediate state between the more differentiated Mit group and dedifferentiated EC groups. This group share features of epithelioid and stromal-rich areas within the tumor.

We found an association between the patients in the group consisting of the EC, EC-Mit, and Mit subtypes and disease stage IV (Fisher exact test, FDR = 0.012, Table S3B). Patients with tumors in the EC, EC-Mit, and Mit subtypes had an increased risk of developing distant metastases and shorter survival times from the detection of the first metastasis. They were associated with overall survival (OS) of less than five years when compared to patients with metastases belonging to the Mit-Im and EC-Im subtypes (Fisher exact test, FDR < 0.03). Other significant associations between the subtypes and histopathological and clinical parameters can be found in Table S3B. In addition, as expected, Kaplan-Meier analysis showed that patients within the immune subtypes (Mit-Im + EC-Im) had a significantly better prognosis than patients with metastases belonging to other subtypes (EC + EC-Mit + Mit) (Figure 1E). A similar trend was observed in the survival analysis of these two groups when considering patients in stage IIIB and IIIC, respectively (Figure S2C and S2D).

Interestingly, there were no significant associations between the proteomic subtypes and the presence of BRAF V600E or NRAS Q61K/R mutations, which indicates that the subtypes are not driven by the main mutational events occurring in melanoma, instead providing an orthogonal classification. Also, several metastases from patients in disease stage IIIA/IIIB to IIIB clustered with metastases from patients in stage IV across the more aggressive subtypes EC, EC-Mit, and Mit (Figure 1A). Thus, the proteomic subtypes classify melanoma lymph node metastases beyond the level of clinical staging and genomic driver mutations.

### Expression of melanoma and phenotype switching markers across the subtypes

Protein expression levels of known melanoma markers ^26^, such as MITF, MLANA, PMEL, TYR, and SOX10, had the lowest expression in EC-Mit samples (ANOVA, FDR < 6·10^-4^) and had a significantly lower expression in the EC-like subtypes when compared to the Mit-like subtypes (Figure 2A). MITF is one of the key regulators of melanoma differentiation and has been implicated in playing a role in dedifferentiation, commonly referred to as melanoma phenotype switching. MITF expression has been associated with survival, cell cycle control, invasion, senescence, and DNA damage repair ^27,28^. Higher expression of MLANA and TYR have also been linked to phenotype switching in melanoma and is associated with a more differentiated phenotype ^29–31^. Transcript and protein levels of these genes were higher in the Mit and the Mit-Im compared with the other subtypes (Figures 2A and 2B).

**Figure 2.**
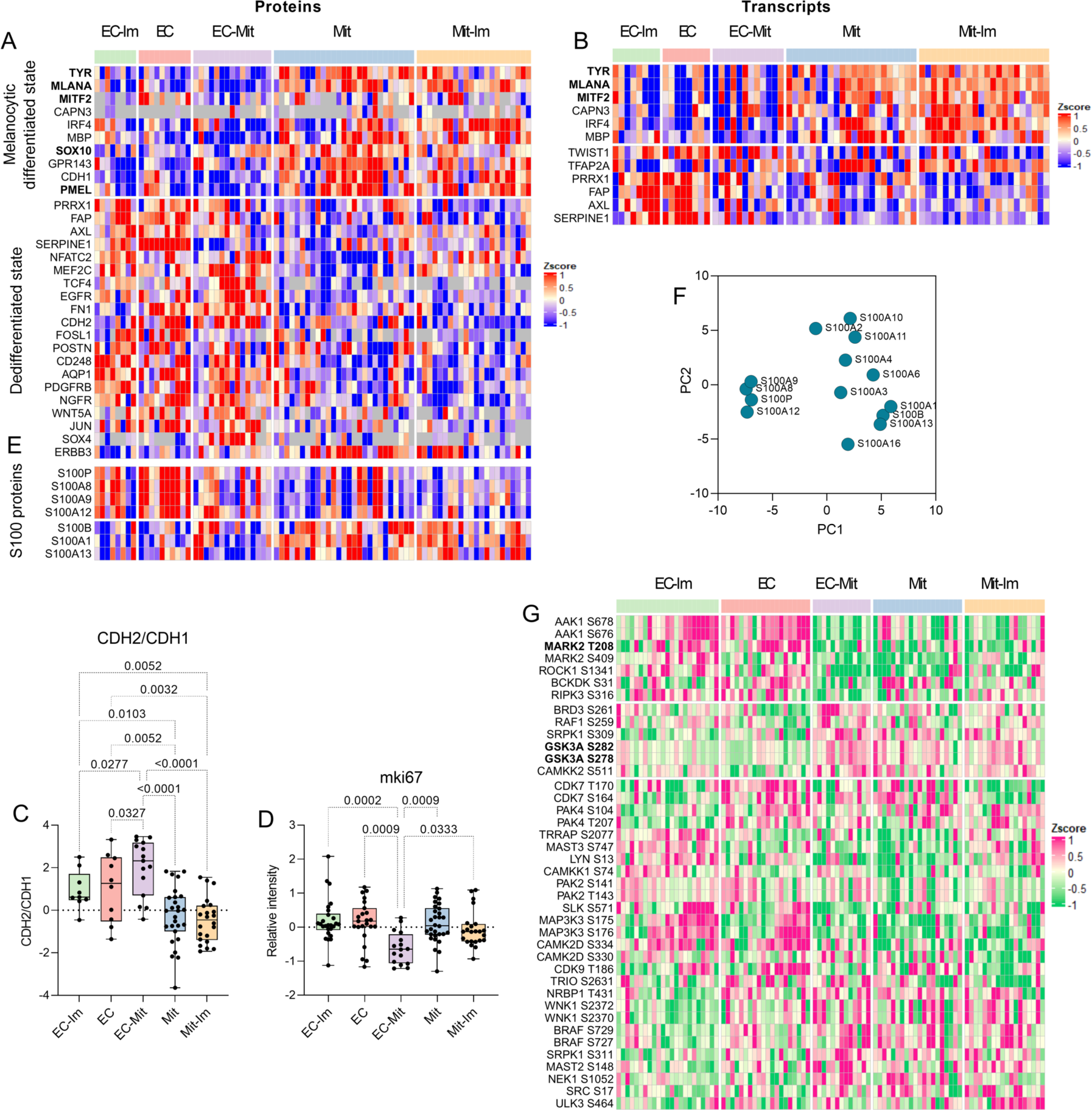
Markers of melanoma and various phenotypic states among the five proteomic subtypes. (A) Protein expression of melanoma markers and markers of EMT in each subtype. In bold, melanoma markers commonly used in clinical practice using IHC. (B) Transcript expression of melanoma markers and markers of EMT in each subtype. In bold, melanoma markers commonly used in clinical practice using IHC. (C) Ratio comparison of the EMT markers CDH1 and CDH2 across the proteomic subtypes. (D) Protein expression of the proliferation marker mki67 across the proteomic subtypes. (E) S100 protein expression across the proteomic subtypes. (F) Principal component analysis of the S100 protein expression. (G) Dysregulated, activated kinases across the proteomic subtypes

The protein expression of the EMT hallmark genes, CDH1 (E-Cadherin) and CDH2 (N-Cadherin) supported the phenotype-switching traits observed at the histological level. We found these proteins to be differentially expressed among the subtypes resulting in a higher relative abundance relationship of CDH2 and CDH1 (CDH2/CDH1 ratio) in the EC, EC-Im, and EC-Mit subtypes (ANOVA, adjusted p-value<0.05) compared to the Mit and Mit-Im metastases, indicative of a switched gene expression from CDH1 to CDH2 (Figure 2C and Figure S2E and S2F). The protein and transcript levels of known markers of melanoma phenotype-switching ^30^ were also analyzed. The gene expression of FAP, ERBB3, FOSL1, and SERPINE1, which have been associated with a dedifferentiated state, were significantly higher in the EC or EC-Im compared with the Mit and Mit-Im subtypes. Also, protein levels of SOX10 and GPR143, associated with a more differentiated state, were significantly higher in the Mit and Mit-Im, compared to the EC or EC-Im subtypes. Moreover, there was a significant differential expression of the SOX4, FOSL1, and SERPINE1 markers between the EC and EC-Mit proteomic subtypes. There was no significant expression difference for VIM, SNAI1/2, and ZEB1/2 transcription factors among the metastatic subtypes, which might reflect the peculiarities of the various phenotypes and differentiation states in heterogeneous metastases.

Overall, the analysis supports a more differentiated state for the Mit and Mit-Im subtypes and a dedifferentiated state for metastases of the EC, EC-Im, and EC-Mit subtypes, matching the histological observations. The presence of several dedifferentiated states is consistent with the increasing evidence that phenotypic transitions encompass more complex dynamics than the extremely differentiated and dedifferentiated states ^32^

The EC-Mit metastases showed significantly lower expression of the proliferation marker Ki67 compared to the other subtypes, which is suggestive of a more invasive phenotype. Interestingly, there was no significant difference in proliferation among the EC and EC-Im subtypes compared to the more differentiated Mit and Mit-Im subtypes (Figure 2D). This challenges the assumed association between invasiveness and proliferation, pointing to a non-exclusive behavior of these features as has been highlighted in previous studies ^33^.

### Selective and coordinated regulation of S100 family members

S100 antigenicity is used to identify poorly differentiated metastatic melanoma, and this protein family has been implicated in cell proliferation, metastasis, angiogenesis, invasion, and inflammation ^26,34–39^. Selected members of the S100 protein family were divided into two groups (Figures 2E and 2F). The first group, consisting of S100P, S100A8, S100A9, and S100A12, was significantly upregulated in the EC subtypes (FDR < 0.003). S100P and S10012 have a diagnostic and prognostic value in many human cancers ^37,40^, while the contribution of S100A8 and S100A9 to melanoma biology is not fully understood. In the second group, S100B, S100A1, and S100A13 were significantly upregulated in both mitochondrial subtypes compared to the EC subtypes, with FDR = 0.0085 and p-value = 0.031, respectively. S100B inhibits p53 phosphorylation and is a serum biomarker of advanced disease stage, poor therapeutic response, and low patient survival ^37,41,42^. S100A1 is known for dysregulating proliferation ^38^ and interacting with the mitochondria ^43^. Overall, the results suggest a selective and coordinated regulation for certain S100 family members within the different proteomic subtypes, which, together with other melanoma markers, may be used to discriminate between proteomic subtypes with poor prognosis.

### Expression of targetable phosphorylated kinases among the subtypes

To identify potential therapeutic targets specific to each proteomic subtype, the phosphoproteomic data on kinases were used as “potential kinase activation surrogates” ^44–47^. We found five phosphosites significantly upregulated in kinases in the EC-like subtypes, including AAK1, MARK2, ROCK1, BCKDK, and RIPK3 (ANOVA p-value < 0.05) (Figure 2G). Phosphorylation at Thr208 of MARK2 was located in the activation loop, which may lead to loss of cell polarity through interfering with microtubule stability and to an unfavorable prognosis, which has been observed in other cancers ^48^ (Figure S2G). In the EC-Mit, Mit, and Mit-Im subtypes, we found upregulation of phosphosites in the kinases BRD3, RAF1, SRPK1, GSK3A, and CMKK2. In GSK3A, the phosphorylated Ser278 and Ser282 are located in the activation loop, flanking the known activation site Tyr279 (Figure S2H). GSK3A was recently linked to cancer stem cells and drug resistance ^49^. Decreasing GSK3A protein levels using siRNA and pharmacological targeting reduced melanoma tumor development in murine models ^50^. Thus, GSK3A could be a promising target for subtype-specific treatment in melanoma.

### A molecular and pathway-level understanding of melanoma histopathological features

To identify relationships between clinical and histopathological features and biological pathways across omics datasets, we used ICA combined with gene set enrichment analysis (GSEA) (Table S4A-D). Variables such as tumor cell content, adjacent lymph node tissue, and necrosis contents were significantly related to many Reactome pathways supported simultaneously by proteomic, phosphoproteomic, and transcriptomic data (Table S4D). Furthermore, the association of one or several features to particular ICs exhibited direct or reverse relationships towards one another (Figure 3A and 3B). For example, we found that proteomic IC 48 was positively associated with closely related parameters such as adjacent lymph node, lymphocyte density, and lymphocytic score and was negatively associated with tumor content. Similar associations were observed in ICs 10 and 92 for the phosphoproteomic and transcriptomic datasets, respectively. These ICs shared enrichment in immune-related pathways such as TCR signaling, PD-1 signaling, IFN signaling, neutrophil degranulation, and complement and coagulation cascades. In addition, specific pathways attributed to protein, transcript, and phosphoprotein expression were found, including apoptosis, antigen processing, and presentation and signaling by Rho GTPases, respectively. The latter has also been associated with triggering multiple immune functions ^51^.

**Figure 3.**
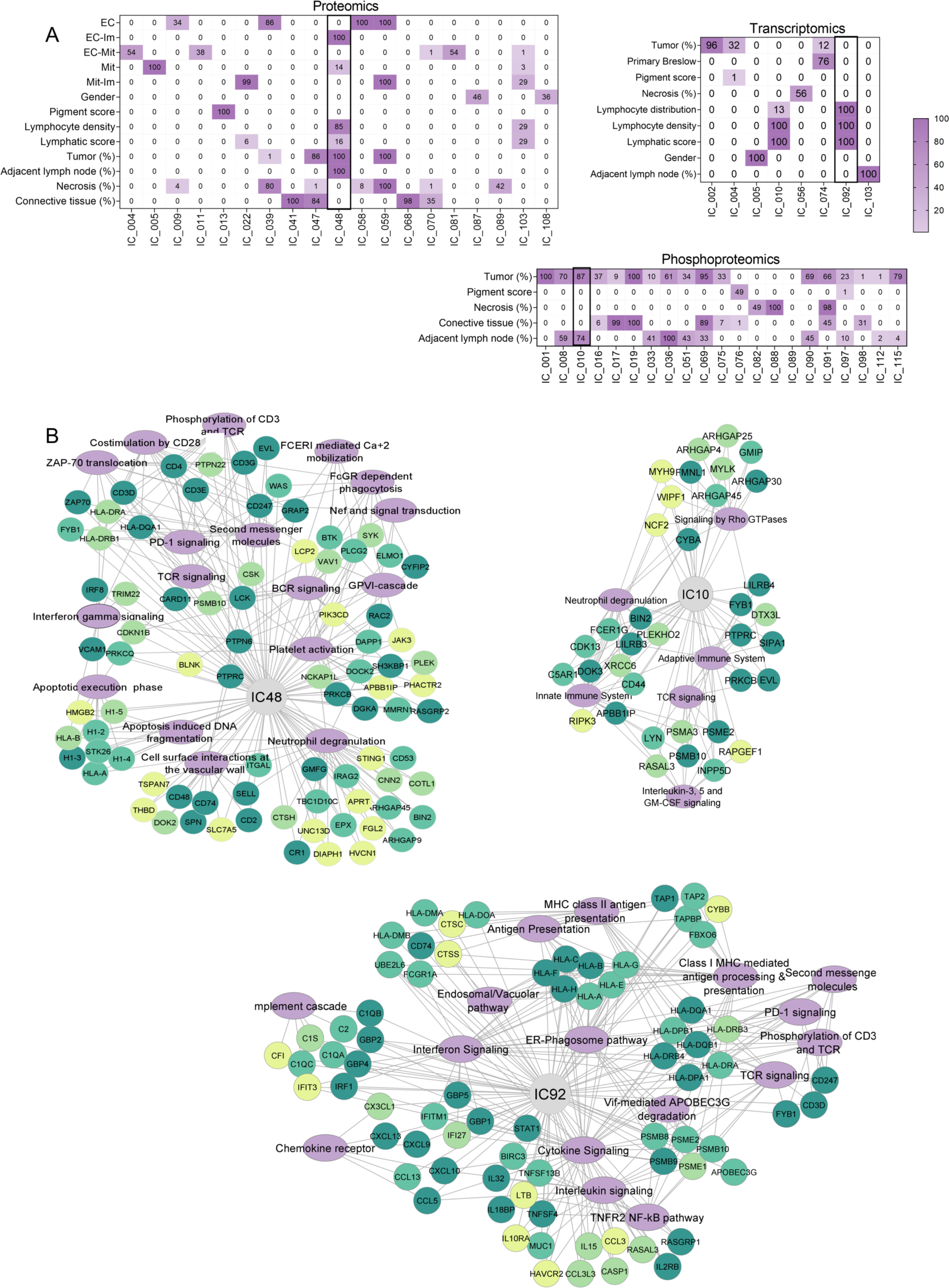
ICA connects pathways with clinical and histopathological features. (A) Percentage of significant correlations between independent components (ICs) and clinical and histological features (p-value < 0.00001) for each omics dataset. IC showing the highest overall correlation percentage in each dataset is highlighted with black contour. (B) Interconnections between proteins with an IC score > 2 and pathways based on enrichment analysis using the Reactome database (FDR < 0.05) for the ICs 48 (proteomics), 10 (phosphoproteomics), and 92 (transcriptomics). The green color scale of the protein nodes indicates their quartile (dark green corresponds to the highest protein score).

### Proteomic signatures and patient survival stratify BRAF V600 positive metastases

A previous study of 16 patients with metastatic melanoma quantified the level of BRAF V600E mutated protein ^17^. Higher expression was associated with lower immune response and increased cell proliferation, leading to tumor progression and shorter survival. We used INGRID to investigate whether these results could be extrapolated to a cohort of 49 metastases of which 67% lacked data on BRAF mutation expression (Figure S3A and STAR Methods). The analysis resulted in a classification of patients into low, medium, and high mortality risk groups. Patients with different mortality risks were identified and classified into low, medium, and high mortality risk groups (Figure 4A). Notably, eight out of nine patients with high BRAF V600E mutated protein levels ended up in the high and medium-risk groups (Figure 4B). In contrast, most patients with low levels of BRAF V600E mutation had a low mortality risk.

**Figure 4.**
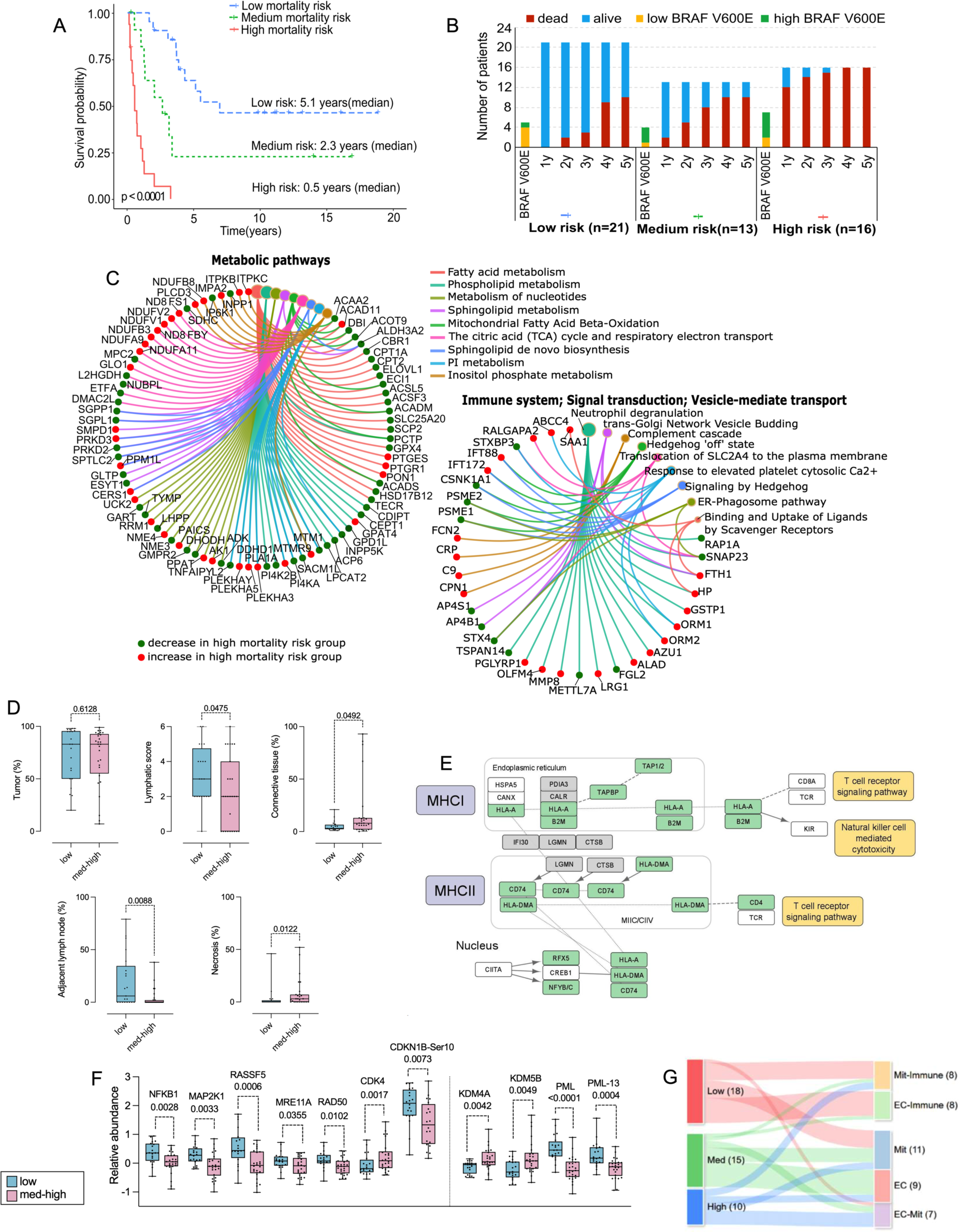
Insights from studying BRAF V600E mutated metastases. (A) Kaplan-Meier curves of BRAF V600 subgroups of patients. Subgroups colored by survival probabilities: red (high risk of mortality, n = 16), green (medium risk, n = 12), and blue (low risk, n = 21). Median survival times for the three groups are shown. (B) Status (alive/dead) distribution of BRAF patients after 1 to 5 years from sample collection. X-axis: patient distribution stratified according to mortality risk classification. The number of patients in which levels of mutated BRAF V600E protein could be quantified (Betancourt et al., 2019b) is indicated in yellow (low) and green (high). (C) Proteins from the most significant pathways enriched in the patients with BRAF mutation. Green and red indicate the decrease and increase of protein expression, respectively, in the high mortality risk group. (D) Histological features linked to BRAF mortality groups. (E) Significantly upregulated proteins (green) of the antigen processing and presentation pathway in the low-mortality risk group compared to the med-high risk group. Identified proteins are shown in (grey). (F) Proteins and phosphorylation sites linked to cellular senescence and their expression patterns between the low- and med-high mortality risk groups. (G) Association between BRAF mortality groups and the five proteomic subtypes.

The proteins involved in classifying these risk groups were mainly enriched in pathways related to metabolism, the immune system, and signal transduction (Figure 4C, Table S5A). Neutrophil degranulation, the complement cascade, mRNA transcription, TGF-beta signaling, and DNA repair were positively related to high mortality risk. On the other hand, signal transduction, vesicle-mediate transport, mitochondrial fatty acid beta-oxidation, fatty acid metabolism, and nucleotide metabolism were positively related to low mortality risk. We also investigated whether disparities between BRAF patients with low and medium-high mortality risks were apparent among the histological features. There were no significant differences in tumor cell content between the low and medium-high mortality risk groups (Figure 4D). However, low-risk metastases, as opposed to the medium-high risk group, were characterized by higher lymphatic scores (ANOVA test, FDR < 0.05) and adjacent lymph node content (Kruskal-Wallis test, FDR < 0.05) as well as lower tumor-derived connective tissue and necrosis content (Kruskal-Wallis test, FDR < 0.05).

In a previous study, we observed a tendency towards larger cells with multiple nuclei in the metastases of the low BRAF V600 group. This feature has been associated with cellular senescence ^52–54^. Transcriptomic studies with cultured melanoma cells and mouse models have found a link between oncogene-induced senescence (OIS) and an increase in the MHC II molecules, which is known to be associated with a favorable disease outcome ^55,56^. In the low-mortality risk group, in agreement with the higher lymphatic scores assigned to these tumors, we found an upregulation of most proteins related to antigen processing and presentation by MHC II (Figure 4E). This proposes an OIS-like phenotype that could act as a tumor suppressor and lead to the better outcome observed in this group.

Enrichment of the differentially expressed proteins and phosphosites between the two mortality risk groups showed an upregulation of interferon-gamma response in the low-mortality risk group (Figure S3C) which may also induce senescence and activate MHC I antigen presentation ^57,58^. Indeed, we found an upregulation of many proteins within the MHC I pathway in the low-mortality risk group (Figure 4E). This suggests an increased susceptibility to cell-mediated cytotoxicity in this group and a possible contribution to the OIS-like phenotype, resulting in a better prognosis ^59^.

Several proteins from the KEGG cellular senescence pathway, such NFKB1, MAP2K1, RASSSF5, MRE11A, and RAD50, were upregulated in the low-mortality risk group, whereas CDK4, which, when bound to Cyclin D inhibits Rb, was downregulated (Figure 4F). Moreover, a significantly increased phosphorylation of CDKN1B at Ser-10 was observed (Figure 4F). This is the most important phosphorylation site in resting cells as it inhibits CDK2 activity, prohibiting G1 to S phase cell cycle transition.

To further support the OIS-like state in the low-mortality risk group, JmjC demethylases such as KDM4A and 5B, known to regulate senescence directly or indirectly, were found to be downregulated in these tumors (Figure 4F) ^60^. KDM5B has been shown to endorse the demethylation of H3K4me3/2, resulting in the silencing of E2F target gene promoters through direct interaction with Rb ^61,62^. The demethylase activity of KDM5B is crucial for DNA damage repair and genomic instability, whereby the inhibition of KDM5B leads to the activation of p53, inhibiting cell proliferation ^63–65^. The downregulation of KDM4A and KDM5B ^63–65^ is known to activate the p53 pathway, triggering senescence and knockdown of KDM4A that can lead to the buildup of promyelocytic leukemia (PML) nuclear bodies, a marker of senescence, also upregulated in the low-mortality risk group (Figure 4F) ^66^. Regulation of the PML protein, a known tumor suppressor ^67^, largely depends on several phosphorylations of the protein. We found an upregulation of phosphorylation at Ser-8 and Ser-36, which increases PLM protein accumulation, and at Ser-403 and Ser-518, which promotes PML protein degradation in the BRAF low mortality risk group, suggesting an increased protein turnover (Figure S3C) ^68–71^. KDM4A and KDM5B can be considered proto-oncogenes, and targeting these demethylases could potentially result in tumor suppression, which makes them attractive therapeutic targets in melanoma.

A significant association was found between the low-risk mortality BRAF subgroup and the proteomic subtypes with better prognosis (EC-Im and Mit-Im) (Fisher exact test, FDR=0.002). In comparison, the medium and high-risk BRAF subgroups were significantly associated (Fisher exact test FDR=0.002) with the subtypes having worse prognoses (EC, Mit, and EC-Mit) (Figure 4G). The medium-high BRAF V600 mortality risk group was also associated with, an increase risk of developing distant metastases, shorter survival from the detection of the first metastasis, and shorter overall survival (Table S5B).

The differences in protein profiles, biological pathway enrichments, and histological features support a heterogeneous BRAF mutation expression. This is defined by two major groups of BRAF V600 positive metastases linked to the escape from or exposure to immune surveillance. For the latter, we propose an OIS-like phenotype as a tumor-suppressing mechanism in patients with lower mortality risks, contributing to their better prognosis.

### Melanoma-associated single amino acid variants (SAAVs)

By further exploring the melanoma proteome, we identified 1,015 SAAVs in 828 proteins in the lymph node metastases (Figure S4A, Table S6A, STAR Methods). Interrogation of the CanProVar database and the Cancer Gene Census resulted in 27 and 24 SAAVs related to cancer or belonging to genes with mutations implicated in cancer, respectively (Table S6A). In addition, 30 SAAVs were predicted to be cancer-promoting by FATHMM ^72,73^.

We matched identified proteins with SAAVs to signaling pathways and biological processes recurrently dysregulated in melanoma (Figure S4B) ^74^. The largest number of SAAVs was found in the PI3K/AKT and MAPK signaling pathways. Besides the well-known role of mutations in members of the MAPK pathways such as BRAF and NRAS, an increasing number of studies have linked gene polymorphism and genetic variants to the members of the PI3K/AKT signaling pathway as susceptibility for cancer development ^75–77^. Most of the SAAV-affecting proteins of the PI3K/AKT signaling pathway were structurally or functionally associated with the ECM. Indeed, Reactome and KEGG pathway analyses of proteins with SAAVs showed enrichment in three general categories, with ECM-related processes as the most enriched pathway. This was followed by cellular metabolism and the complement and coagulation cascade (Figure S4C and S4D). Less than 2% (19) of all the SAAVs, originated from samples with <50% tumor content, indicating that the ECM-related protein variants are likely produced by the melanoma cells. This finding is in line with a recent Pan-Cancer genomic study, which revealed that a higher copy number and more missense mutational alterations are present in the ECM genes compared with the rest of the genome ^78^.

Quantitative analysis of the SAAVs expression resulted in 52 differentially expressed variants across the proteomic subtypes (ANOVA, FDR<0.05), where the highest number of overexpressed SAAVs was found in the EC subtype (Figure 5A). These SAAVs underlined an active role in the remodeling of the ECM and dysregulation of the complement and coagulation pathway, processes that are likely associated with phenotype switching of melanoma cells. This is in agreement with the enriched pathway results for all identified variants.

**Figure 5.**
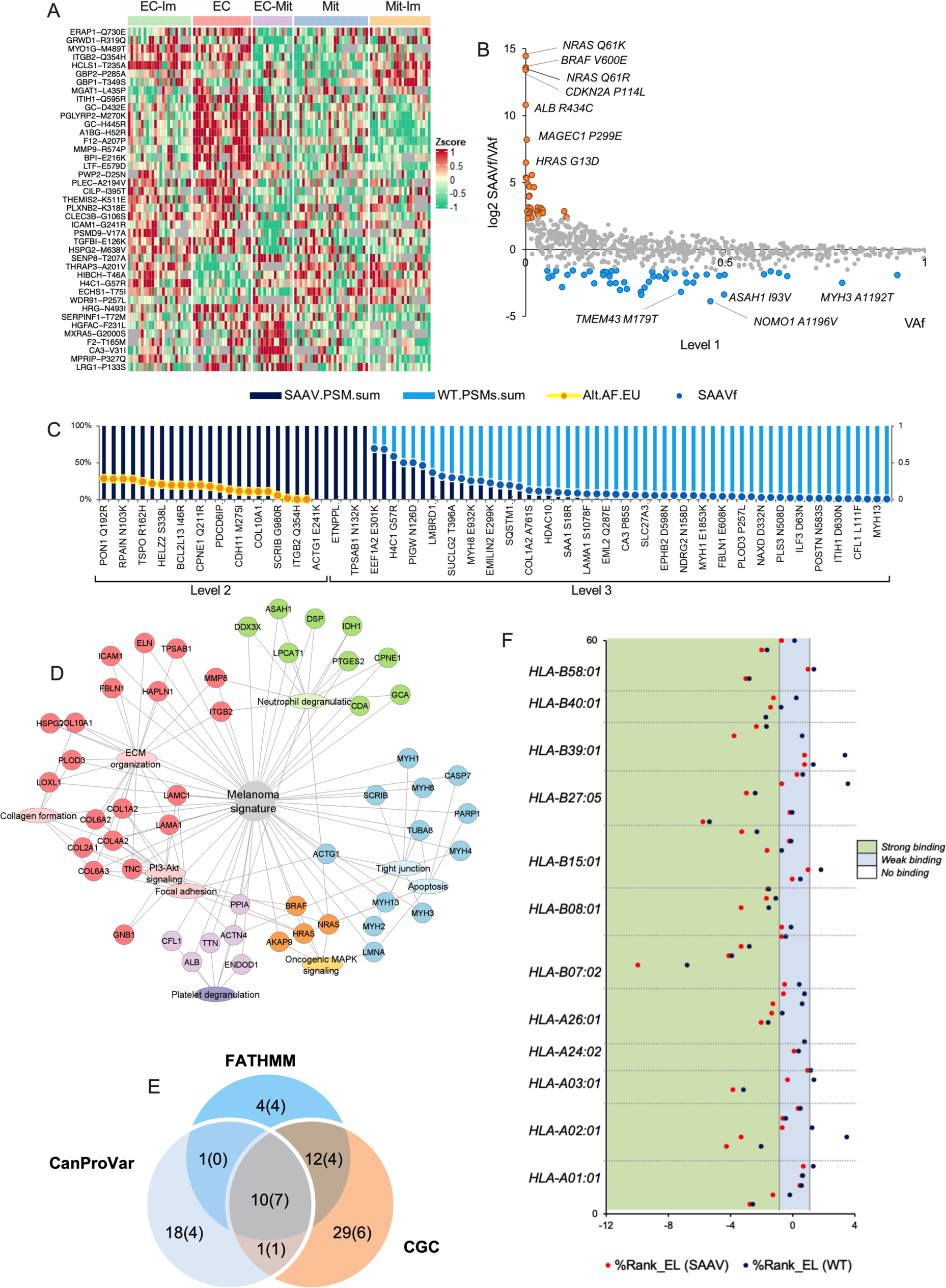
The landscape of SAAVs in melanoma. (A) Differentially expressed SAAVs across the proteomic subtypes. (B) Level 1 of the melanoma-associated SAAV signature defined in this study, with over-(orange) and under-(blue) represented SAAVs from what is expected based on SAAVf and Vaf ratio. (C) Proteomic evidence (#PSMs of wild-type and SAAV peptides) and genomic data (VAf) for Levels 2 and 3 of melanoma-associated SAAV signature defined in this study. (D) Melanoma-associated SAAV signature interconnections between corresponding proteins/genes and pathways, based on enrichment analysis using KEGG and the Reactome databases. (E) Overlap of the SAAVs and genes annotated by CanProvar and CGS or predicted by FATHMM as implicated in cancer. The numbers in parenthesis represent the melanoma-associated SAAV signature covered by these predictors. (F) Predicted affinity SAAV-neoantigen candidates per HLA, which ranked better than wild-type counterparts using NetMHC.

To predict which SAAVs are significantly related to melanoma we searched the dbSNP Short Genetic Variations database ^79^ for the European population’s corresponding allele frequencies (VAf). As a result, we generated a signature of 167 melanoma-associated SAAVs classified into levels 1, 2, and 3, according to genomic data and proteomic evidence (Figure 5B and 5C and Table 6A) (STAR METHODS). These SAAVs displayed the same pathway enrichment as described above (Figure 5D). The signature captured 26 critical SAAVs among the genes and mutations reported by CanProVar and The Cancer Gene Census or predicted by FATHMM (Figure 5E). This included the melanoma driver mutations NRAS Q61K/R, BRAF V600E, CDKN2A P114L, and HRAS G13 (Figure 5C). These findings served as evidence to support the proposed signature of melanoma-associated SAAVs. Interestingly, we also found five SAAVs in heavy chains of muscle myosin II complex (MYH1 E1853K, MYH2 E486K, MYH4 N1627I, MYH8 E932K, and MYH13 D1765N), which presented the highest number of variants originated from the same loci 17p13.1 in our dataset (Table S6A). This is the same loci of TP53, a frequently mutated tumor suppressor credited as a hallmark of cancer. Myosin II is required for cell contractility, cytoskeleton reorganization, and cytokinesis. Different reports associated mutations and polymorphism in heavy chains of muscle myosin II to cancer predisposition ^80^ and non-cancer-related diseases ^81,82^. We hypothesize that these variants are reactivated by melanoma cells as part of cytoskeletal remodeling by myosins ^83^.

Tumor mutational burden can predict response to immunotherapy in melanoma ^84^, which suggests that mutated peptides binding to MHC I class molecules can be the primary origin of target antigens. To investigate neoantigens with the potential to express identified SAAVs, we aligned the amino acid sequence of the SAAV-bearing peptides to a large experimental dataset of a melanoma-associated immunopeptidome ^85^. This resulted in 56 SAAVs that could be presented as variant peptide ligands by the HLA I complex (Table S6A). According to the NetMHC prediction tool ^86^, 61 SAAV-altered peptides ranked better as HLA class I neoepitopes than their wild-type counterparts (Table S6B). This included 15 cases where the latter fell out of the specified threshold (%Rank_EL < 2) for binding prediction (Figure 5F and Table S6B). The variant peptide ligands CYB5R1 N44S, CD300LF Q218R, GCA S80A, and QARS N285S were the best candidates for the HLA allotypes HLA-A26:01, HLA-B07:02, HLA-B15:01, and HLA-B58:01, respectively. Among the significant HLA I peptide ligand variants, we found CDKN2A P114L, ARL6IP6 R56L, GCA S80A, LOXL1 R141L, CFL1 L111F, HSPB1 I179N, and TGFBI E126K, which are a part of the melanoma-associated SAAV signature defined in this study. The results indicated that knowledge of variant expression, supported by melanoma-associated immunopeptidome and MHC I binding prediction tools, may lead to the discovery of neoantigen candidates as targets of anti-tumor immune responses.

### The tumor microenvironment is a key player in disease progression

Differences in the content of adjacent lymph nodes and the tumor-derived connective tissue between the BRAF V600 mortality risk groups and among the proteomic subtypes pointed to a link between the TME and patient outcome. Therefore, a subset of 29 samples with less than 50% tumor cells was analyzed. Univariate receiver operating characteristic (ROC) curve analysis separated groups of patients with differences in survival (DSS< or ≥3 years) (Figure 6A). The groups were named high or low lymph node (HLN or LLN) and high or low connective tissue (HCT or LCT) based on the content of these histological features in the tumors. With minor differences, the HLN and LCT or LLN and HCT groups allocated the same patient subsets (Figure 6B and 6C), reflecting the significant correlation (Spearman - 0.84, p-value=1.64E-08) and complementarity between LN and CT contents in this subset.

**Figure 6.**
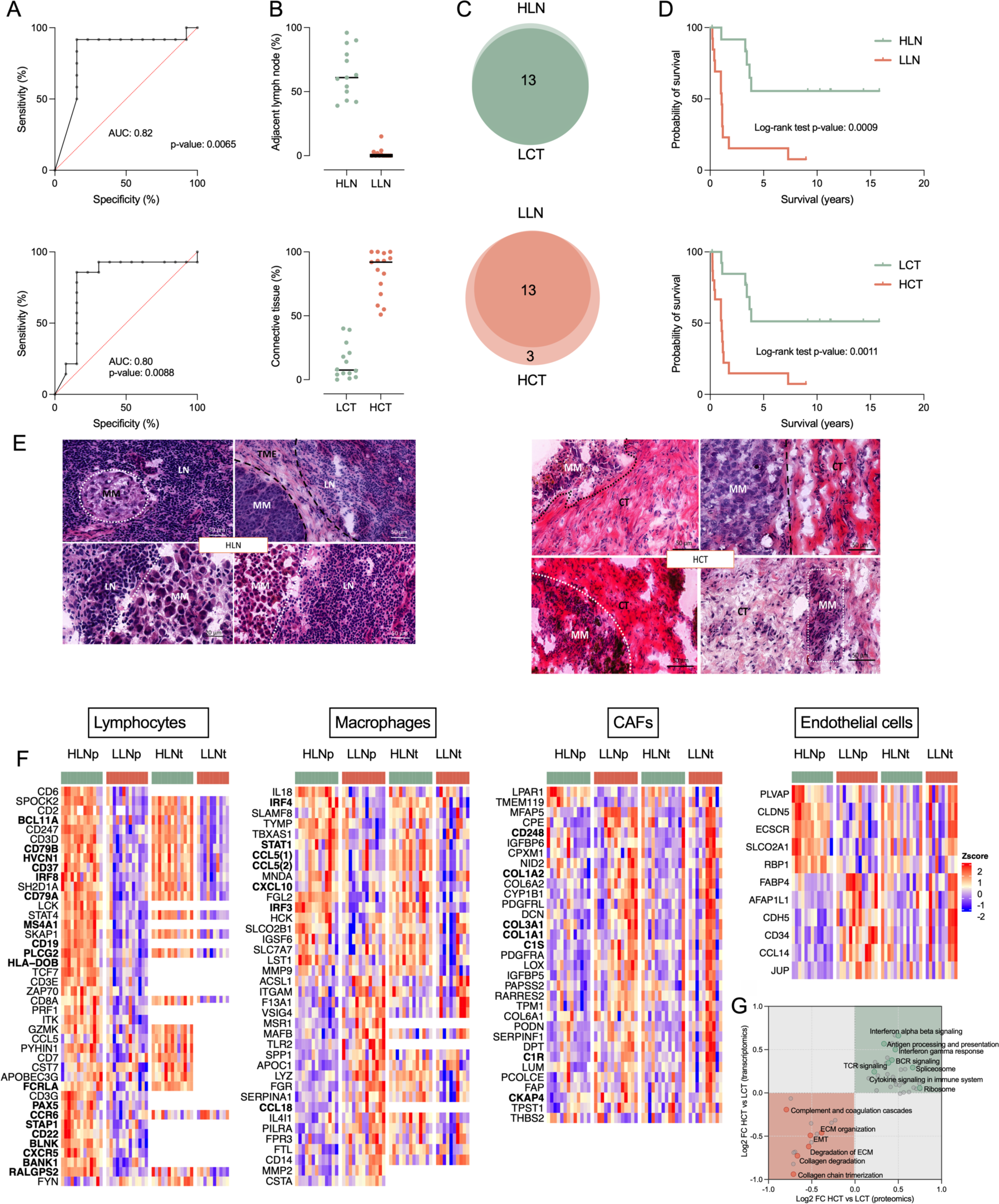
The tumor microenvironment composition is an independent prognostic marker in melanoma lymph node metastases. (A) ROCs of tumor-associated adjacent lymph node (top) and connective tissue (bottom) for DSS< or ≥3 years. (B) Adjacent lymph node (top, cut-off = 27%) and connective tissue (bottom, cut-off = 45.5%) content in the TME for the subgroups of samples generated from the ROC analysis. (C) Venn diagram of the patient overlap between the HLN and LCT groups and between the LLN and HCT groups. (D) DSS probability for patients with tumors grouped based on their content, HLN and LLN (top) or HCT and LCT (bottom). (E) Histological images from different tumor areas in the HLN and HCT groups. (F) Cell-specific signatures at protein (HLN.P and LLN.P) and transcript level (HLN.T and LLN.T) for the HLN and LLN groups. The displayed markers were significant in either the proteomic or transcriptomic analyses (t-test p-value < 0.05). Bold indicates significance in both. (G) 2D enrichment analysis displaying significant pathways (FDR < 0.001) commonly dysregulated on the proteomic and transcriptomic levels between the HLN and LLN groups.

Patients within the HLN and LCT groups had a better prognosis, while the opposite was found for patients in the LLN and HCT groups (Figure 6D). Strikingly, Cox regression models showed that both LN (Cox coefficient = -1.661, p-value = 0.001) and CT (Cox coefficient = 1.718, p-value = 0.001) contents were better indicators of prognosis than disease stage (Cox coefficient=1.265, p-value=0.008). In addition, multivariate Cox regression analysis considering age, gender, and disease stage showed an increased risk of developing distant metastasis for the LLN group compared to the HLN group, with a Hazard ratio (HR) of 5.96 (95% CI: 1.629 to 28.96, p-value=0.021). A similar analysis based on OS showed that patients with metastases in the LLN group have shorter life spans than those in the HLN group (HR= 19.91, 95% CI: 3.350 to 173.4, p-value=0.0023).

The TME of the HLN (also LCT) group was dominated by immune cells, both surrounding and infiltrating the tumor. On the contrary, the LLN (also HCT) group displayed large areas of stroma and fatty tissue mixed with tumor cells (Figure 6E). These observations were supported by cell-specific transcript signatures based on a single-cell study of melanoma TME ^32,87^, which were confirmed at the protein level in our study (Figure 6F). The HLN group was enriched in markers mainly expressed by B and T lymphocytes, macrophages, and endothelial cells (Figure 6F). The enriched pathways were related to antigen processing and presentation, ribosome, and B and T cell receptor signaling at both proteomic and transcriptomic levels (Figure 6G and Table S7A-B). On the contrary, the LLN group displayed enrichment in markers of cancer-associated fibroblasts (CAFs), macrophages, and endothelial cells. Markers of CAFs were highly upregulated, such as collagens, complement components, and growth factor proteins. This group was also enriched in pathways related to the complement and coagulation cascades, EMT, ECM organization, and collagen chain trimerization and degradation. Dysregulated collagen synthesis and assembly are common driving factors in many cancers ^88^.

Among the upregulated proteins in the LLN group, LOX (lysyl oxidase enzymes), which catalyze the crosslinking of collagens and elastin, increasing the tissue stiffness, has been proven to promote tumor progression through increased integrin signaling and EMT ^88^. In addition, upregulation in the fibroblast activation protein (FAP), which is known to be expressed in tumor mesenchymal and epithelial cells, enhances migration and invasion ^89–91^. Consequently, the TME and its cell composition influence the disease progression fundamentally. This is in line with previous studies in breast and colon cancer, where the tumor-stroma ratio in lymph node tissue was linked to prognosis, and a high stromal score (>50%) was associated with a more aggressive tumor progression ^92–94^. The results fuel the need to explore such variables in larger cohorts since the histopathological assessment could be readily implemented in clinical practice and used for better-informed medical decisions.

### Multi-omics data pinpoint proteins related to patient survival

To find biomarkers related to patient survival, we performed two different analyses. First, outlier analysis (STAR methods), which is based on survival groups (0.5, 1, 3, >5 years) was utilized. For each of these survival groups, we selected the enriched transcript-, protein- and phosphosite outliers (Table S8A). Secondly, a regularized Cox regression (STAR methods) was performed to complement with additional markers of survival (Table S8B). As a result, we found 103 proteins, 44 transcripts, and 21 phosphosites significantly related to survival (outlier FDR < 0.05, Cox score >30).

The proteins ADAM10, HMOX1, FGA, DDX11, SCAI, CTNND1, CDK4, PAEP, and PIK3CB were selected based on the above survival analysis, a comprehensive evaluation (Table S8C) and literature search (STAR Methods). Three proteins (ADAM10, FGA, and HMOX1) were chosen based on specific phosphosites linked to survival. For FGA (Ser364) and HMOX1 (Ser229), the phosphorylation of these specific sites was linked to poor survival, while ADAM10 phosphosite Thr719 was enriched in tumors from patients surviving longer. In addition, high protein levels of CDK4, CTNND1, PAEP, DDX11, SCAI, and PIK3CB in the lymph node metastases were all related to poor prognosis.

While the global proteomic analysis was performed on regional lymph node metastases, to expand our understanding of these markers in the earlier disease presentation of the primary melanoma, we assessed the protein expression in an independent cohort of primary melanomas using immunohistochemistry (IHC). The tissue expression of the examined 9 target proteins was assessed in a tissue microarray (TMA) platform generated from 42 patients (Tables S1B and S9C) with primary treatment-naive melanomas progressing into either locoregional or visceral metastatic disease (STAR Methods). The 9 proteins could be localized within the stroma/TME and the melanoma cells (Figures 7B and S5A-D).

**Figure 7.**
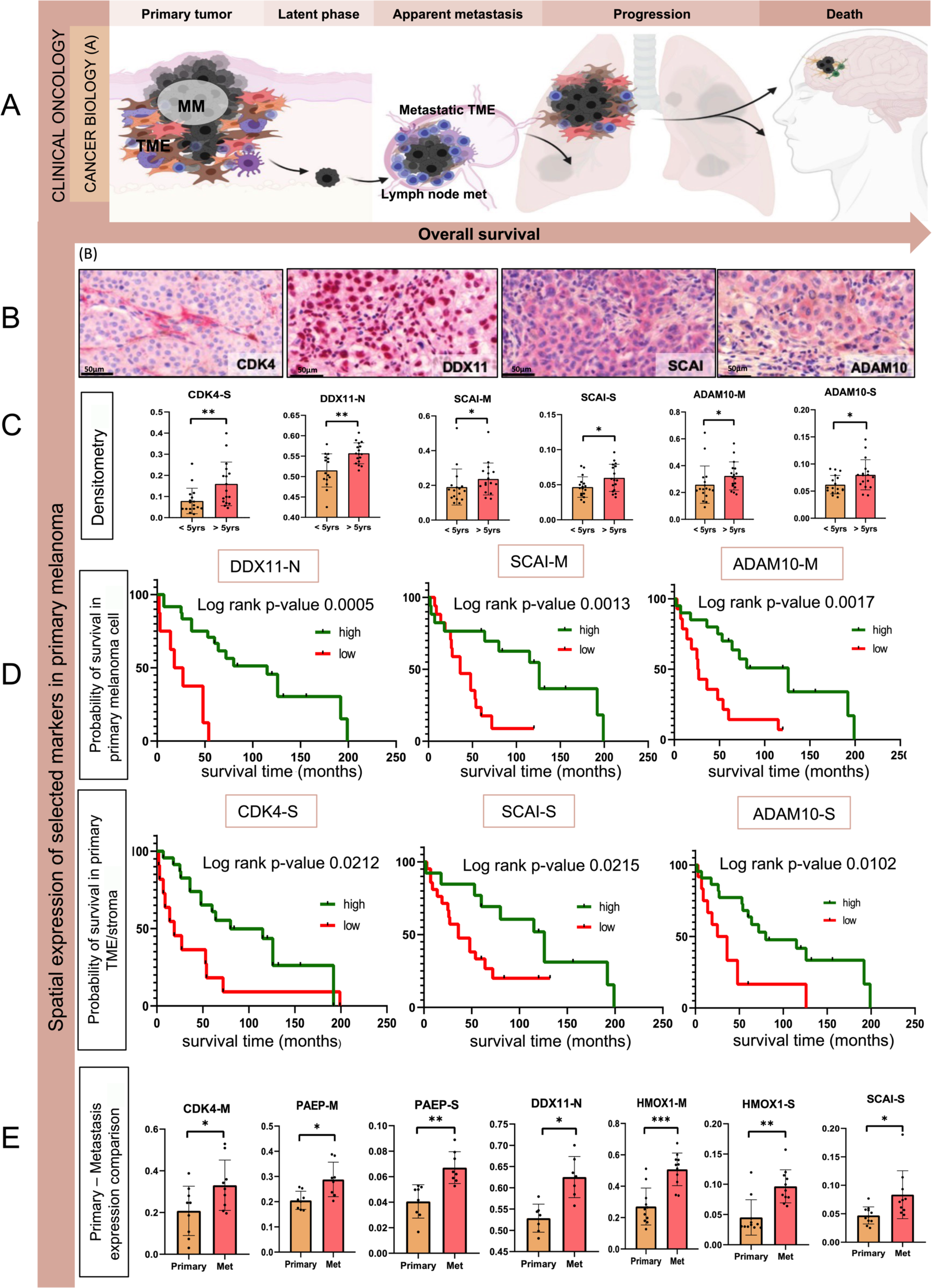
Association of protein marker expression with survival as the disease progress. (A) Visualization of different phases of melanoma showing the relationship between cancer biology and its clinical impact on survival. (B) Representative IHC staining images of markers expression in melanoma and stromal cells of primary tumors. (C) Significant differences found in marker expression associated with OS. (D) Kaplan-Meier survival analyses displaying OS rates for patients in association with high (green) and low (red) expression of the markers in melanoma and stroma cells. (E) Significant differences of the protein markers in pairs of matching primary melanomas and corresponding lymph node metastases.

We found that ADAM10 and SCAI expression was increased in the primary melanomas from patients with OS >5 years compared to tumors from patients with OS <5 years (Mann-Whitney U test p-value<0.05) (Figure 7C and Table S9A). A similar trend was observed in the stromal expression of CDK4 and tumoral DDX11 (p-value<0.05, Figure 7C). In addition, spatial expression-based ROC curve analysis followed by Kaplan-Meier (K-M) analysis revealed that higher ADAM10, FGA, DDX11, PAEP, and SCAI expression in primary melanoma cells could be linked to longer survival, as well as a higher stromal expression for ADAM10, CDK4, SCAI, HMOX1, and PAEP (Figures 7D, S5A-C, and Table S9A).

To follow the dynamics of disease progression, we compared these proteins in up to 9 pairs of matching primary melanomas and lymph node metastases (Table S9A). The expression of most proteins tended to increase from the primary tumor to the corresponding metastasis except for cellular CTNND1 and both cellular and stromal ADAM10 (Figure 7E). In the metastases, CDK4 and DDX11 expression in melanoma cells were significantly higher (Wilcoxon-test p-value < 0.05), and HMOX1 and PAEP expression significantly increased (Wilcoxon-test p-value < 0.05 in both melanoma cells and the stroma/TME), while SCAI expression was significantly increased in the metastatic stroma (Figure 7E).

## DISCUSSION

Melanoma of the skin is heterogeneous regarding cancer cell behavior and the surrounding tumor microenvironment, including the lymphoid, histiocytic, and fibroblastic elements. Cells from the primary melanoma may disseminate early, undetectable, during a latent clinical phase ^95^. Due to the large heterogeneity of melanoma tumors, patient stratification beyond the level of mutations is of utmost importance for improved clinical decisions ^96^. The melanoma samples in the present study were collected prior to the era of immune checkpoint and targeted therapies. Consequently, the molecular and histological characterization of this treatment naïve cohort makes it possible to study the biological characteristics of melanoma metastases in close connection with patient outcomes.

Here we propose proteomic subtypes that integrate microenvironment players such as the immune and stroma components, unlike transcriptomic studies on tumors or cell lines ^7,12,13,97^. This holistic approach enabled refined subtyping of melanoma and significant associations with clinical and histopathological features. For instance, the EC-Im and Mit-Im subtypes were associated with slower metastatic spread and linked with a better prognosis. In contrast, the EC, EC-Mit, and Mit subtypes appeared to escape the immune control and were accompanied by a more aggressive disease. Our results suggest that the subtypes can be distinguished by MS-based abundance of routinely used histopathologic protein markers such as MITF, PMEL (HMB45), MLANA, and certain S100 protein family members ^98^. The S100 proteins appear to have a selective and coordinated regulation across subtypes and in connection with melanoma phenotype switching, which was also observed across the subtype signatures obtained by ICA. These signatures could also serve as diagnostics markers for patient stratification.

The proteomic subtypes unraveled the presence of different states of phenotype-switched melanoma cells, which is in line with prior observations regarding the plasticity and phenotypic diversity of melanoma and contributes to increase knowledge of the pathomorphology within the tumoral tissue ^30,32,99^. Here dedifferentiation appears to be linked to accelerated disease progression due to the association between the EC-Mit subtype and earlier development of distant metastases.

In summary, the proteomic subtypes captures layers of complexity beyond the transcript level by association with molecular, functional, histopathological, and clinical aspects of the disease.

Based on the activation of specific metabolism-, immune system- and signal transduction-related pathways, patients bearing tumors with BRAF V600 mutation could be stratified into subgroups with differing mortality risks. Our results point towards a link between low BRAF V600 expression and a better immune response mediated by a BRAF V600-induced senescence-like phenotype, where the OIS acts as a tumor suppressor through the upregulation of antigen presentation and interferon-gamma signaling, thus improving patient outcomes. The BRAFV600E protein expression appears to have prognostic significance which is independent of the therapy. For metastatic spread, the surrounding niche of TME is necessary for progression ^100^. In a subset of patients with lower tumor cell content, we found a significant association of adjacent lymph node tissue and connective tissue with patient survival, both appearing to be independent prognostic markers in melanoma lymph node metastases. Assessment of tumor-derived connective tissue/tumor or adjacent lymph node tissue/tumor ratio can be accomplished in routine pathology on hematoxylin and eosin-stained slides. It might be helpful as a selection criterion for therapy, as suggested for colorectal cancer and breast cancer ^101–103^. Thus, our study underscores the importance of further investigating these phenomena in larger cohorts to obtain deeper insights into the relationship between the tumor and its microenvironment and its association with clinical outcomes.

The aforementioned classifiers may pose challenges and opportunities from diagnostic and therapeutic perspectives, such as matching subtypes and TME subgroups with response to immune checkpoint treatments or to whether all patients harboring the BRAF V600 mutation benefit from targeted therapy. Moreover, these classifiers highlight the importance of the stroma tumor-infiltrating lymphocytes (sTILs) and their immune signature for survival.

Despite melanoma being the cancer type with the highest mutational burden, the literature is scarce when addressing the protein expression of mutations. Here we present a landscape of SAAVs in often dysregulated pathways affected by driver mutations and of less explored variants in melanoma. We found significant enrichment in the expression of SAAVs of ECM-related genes. By remodeling the ECM, these SAAVs may contribute to TME heterogeneity and impact disease progression, as shown by upregulation in the EC subtype. In addition, we generated a signature of over and under-represented SAAVs in melanoma. Future studies are required to decode the role of these SAAVs in aspects of protein function, melanoma predisposition and progression, and their relation to survival. Additionally, we identified several SAAVs with higher peptide-MHC I binding prediction than previously detected wild-type peptide ligands. This highlights the potential of identifying SAAV expression as a valuable source of antigens eliciting tumor-specific immune responses.

Of the nine putative proteomic biomarkers studied in an independent cohort using IHC, CDK4, ADAM10, SCAI and DDX11 seem promising starting points for further investigation of their role in melanoma routine diagnostics. Besides the IHC expression intensity, the difference in spatial expression in melanoma cells and the stroma/TME demonstrated the complexity of the tumor-stroma relationship.

CDK4 is a well-known cancer target that regulates cell cycle and proliferation. In melanoma, mutations, and dysregulation are commonly seen in CDK4 and proteins in its pathways, and several candidate drugs are in clinical trials ^104–106^. Here, high stromal CDK4 levels in primary tumors were correlated with improved survival, while a significant increase in CDK4 levels was observed in metastatic tumor cells. It has been shown that CDK4/CDK6 inhibitors could have an important role in tumor growth ^107^. Their results suggested that *in vitro,* the CDK4/CDK6 inhibited fibroblasts can induce genotype-dependent tumor cell proliferation and prolonged inhibition of senescent cells in the TME. In agreement with our findings, this study raises some questions about the hidden stromal effect of the CDK4 pathways.

ADAM10 is a metalloproteinase that, by cleaving the ectodomains of transmembrane proteins, has a widespread effect on cancer cells and their stromal counterparts ^108,109^. Our phosphoproteomic data points to a link between the conserved phosphosite in ADAM10 and T719, and improved survival, mirrored by protein expression in the primary melanoma cohort. Analogous to the T735 of ADAM17 phosphorylated by MAP kinases ^110–112^, T719 in ADAM10 may activate the protease.

SCAI is a highly conserved protein that acts on the RhoA–Dia1 pathway to regulate invasive cell migration ^114^. SCAI protein has a debated role in cancer prognostics, correlating with a better outcome in breast and lung cancers ^115^. We found that high expression of SCAI in melanoma cells and the TME was associated with better outcomes in primary melanoma.

The helicase DDX11 has a role in chromatid cohesion, influencing proliferation and melanoma cell survival ^116^. DDX11 expression has been reported upregulated during melanoma progression ^116–119^, which mirrored our proteomic and IHC analysis results.

The study also opens venues for a model building to study drug response-resistance relationships, tumor dormancy, and immune response that may lead to the discovery of new therapeutic targets ^120^. The results suggest that tumor stratification, as outlined here, could offer better prospects for the clinical characterization of tumors that may contribute to more informed medical decisions ^121^.

Our study serves the research and clinical community in multiple ways. For basic scientists, it provides an integrated multi-omics dataset that can be used to generate novel hypotheses. For clinicians, our study offers novel ways to predict prognosis and possibly optimize and customize treatment by stratifying patients using histopathology in combination with a proteogenomic characterization.

## MATERIALS AND METHODS

### Collection of malignant melanoma samples and ethical approval

#### Discovery cohort

A total of 137 patients diagnosed with metastatic melanoma between the years 1975 and 2011, were included in the study. For 80% of patients, it was possible to reclassify the disease stage (Table S1A) based on the 8th edition of the American Joint Committee on Cancer (AJCC8). Here, 29 patients were in stage IIIB, 35 in stage IIIC, 10 were classified as IIIB/IIIC, 4 as IIIC/IIID, and 34 patients as stage IV. In addition, four patients were distributed as stages IIA/IIB (2), IIB (1) and IIIA/IIIB (1). The average age ± standard deviation (range) at diagnosis of metastases was 62.3 ± 13.7 (25–89) years, and the cohort had a preponderance of male individuals (65%). Among patients with available overall survival (OS) information, 50% survived less than 5 years. The samples comprised fresh frozen metastatic tissues from lymph node (126), subcutaneous (7), cutaneous (1), and visceral (3), while for five samples the origin could not be established. Different pieces of the same tumor were analyzed in five cases of lymph node metastases resulting in a total number of 142 analyzed samples. The mutational status was determined for 124 of the metastases, with 49 found mutated at BRAF (92% V600E), 36 with NRAS Q61K/R, and 37 tumors that had wild-type variants for both mutations. Only four patients received targeted BRAF treatment with vemurafenib. The histopathological classification of the primary tumors was the following: Nodular melanoma (NM; 54%), Superficial Spreading Melanoma (SSM; 38%), Acral Lentiginous Melanoma (ALM; 5%), Lentigo Malignant Melanoma (LMM; 1%), and Mucosal (2%), where 100 patients had thicker primary tumors (Breslow > 1 mm). A more detailed description of clinical parameters for every patient is provided in Table S1A. The study was approved by the Regional Ethical Committee at Lund University, Southern Sweden (DNR 191/2007, 101/2013 (BioMEL biobank), 2015/266, and 2015/618). All patients provided written informed consent. The study has been performed in compliance with GDPR.

#### Melanoma FFPE primary and metastatic tumor cohort for IHC analysis

The melanoma samples included in the IHC cohort were collected from the Department of Dermatology and Immunology of the University of Szeged. Patients (n=42, from 2001 to 2020) were selected whose primary melanoma is archived as paraffin-embedded tissue blocks. All primary tumors resulted in loco-regional and/or disseminated disease. The tissue microarrays (TMAs) were made from formalin-fixed paraffin-embedded (FFPE) blocks. Clinical information for the 42 patient samples included gender (24 males and 18 females), age at diagnosis of the primary tumor (mean = 61.7 ± 10.1 years), localization of the primary tumor (trunk = 24, lower limbs = 8, upper limbs = 7, head and neck region = 3) and metastases, long term follow up data: disease-free survival interval (DFS) (mean = 23.7 ± 39.1 months), progression-free survival interval (PFS) (mean = 54.6 ± 50.2 months), overall survival (OS) (mean = 59.2 ± 50.8 months), histological subtypes (SSM, NM, ALM, LMM, etc.), pathological TNM staging (according to AJCC cancer staging system, 8th edition), histological parameters of the tumor (Clark level, Breslow, presence of regression and ulceration) and BRAF status. OS was calculated from the date of diagnosis of the primary melanoma to the date of the last follow-up (Table S1B).

According to the guidelines of Declarations of Helsinki, all patient samples were obtained with the approval of the Hungarian Ministry of Human Resources, Deputy State Secretary for National Chief Medical Officer, Department of Health Administration with the ethical authorization number: MEL-PROTEO-001, 4463-6/2018/EÜIG and the date of approval was 12 March 2018. Based on the MEL-PROTEO-001, 4463-6/2018/EÜIG ethical approval, informed consent was not applicable due to the retrospective anonymized FFPE samples.

### Sample acquisition

#### Discovery cohort

Tissue specimens were snap-frozen, or alternatively put on dry ice, within 30 minutes of collection, most samples were frozen within 15 minutes upon surgery, with a small amount of isopentane in liquid nitrogen. Multiple pieces were collected from most of the tumor specimens. The samples were then stored at -80 ℃ in the Melanoma biobank, BioMEL, Region Skåne, Sweden.

### Histopathological analysis

#### Discovery cohort

Stepwise sectioning of the tissues was performed, and on average, three sections were evaluated for each tumor. Frozen tissue sections were placed on glass slides, stained with hematoxylin and eosin, and then placed in an automated slide scanner system (Zeiss Mirax). The tissue content was then evaluated in terms of tumor cells, necrosis, connective tissue, and adjacent background tissue, and features that could be further captured based on morphology were considered. Necrosis did not affect the quality of data. Only 10 out of 142 samples of the cohort have necrosis >20%, and most of them (6) displayed high tumor content (>60%). Therefore, no samples were excluded based on necrosis.

For deeper evaluation, we assessed the properties of the tumor cells (primary pattern, size) and the infiltration of lymphocytes in the tumor mass (an immunoscore was given representing tumor-infiltrating lymphocytes both in the dimension of intensity and extent). All histological parameters were assessed by a board-certified pathologist. To improve comparability between studies, the methods were adjusted to the at-that-time existing protocol from The Cancer Genome Atlas Network, and their analysis of malignant melanoma (TCGA, Cell 2015). The following pathologic parameters were scored for each case: lymphocyte distribution (0-3, 0 = no lymphocytes within the tissue, 1 = lymphocytes present involving <25% of the tissue cross-sectional area, 2 = lymphocytes present in 25 to 50% of the tissue, 3 = lymphocytes present in >50% of tissue); lymphocyte density (0-3; 0 = absent, 1 = mild, 2 = moderate, 3 = severe). The pathologists were provided with written definitions for each parameter and an illustrated guide. Lymphocyte score is defined as the sum of the lymphocyte distribution and density scores (0-6) were also calculated for each case.”

### Sample preparation for mass spectrometry

#### Protein extraction, digestion and C18 desalting

Protein extraction was performed on sectioned (30 x 10 µm), fresh-frozen melanoma tissues using the Bioruptor plus, model UCD-300 (Dieagenode). In total, 142 melanoma samples were lysed in 100 µL lysis buffer containing 4 M urea and 100 mM ammonium bicarbonate. After a brief vortex, samples were sonicated in the Bioruptor for 40 cycles at 4°C. Each cycle consisted of 15 s at high power and 15 s without sonication. The samples were then centrifuged at 10,000 ×g for 10 min at 4°C. The protein content in the supernatant was determined using a colorimetric micro–BCA Protein Assay kit (Thermo Fisher Scientific).

Urea in-solution protein digestion was performed on the AssayMAP Bravo (Agilent Technologies) micro-chromatography platform using the digestion v2.0 protocol. Protein concentrations were adjusted to 2.5 µg/µL and 100 µg of protein from each sample were reduced with 10 mM DTT for 1 h at room temperature (RT) and sequentially alkylated with 20 mM iodoacetamide for 30 min in the dark at RT ^122^. To decrease the urea concentration, the samples were diluted approximately seven times with 100 mM ammonium bicarbonate.

Digestion was performed in two steps at RT. Proteins were first incubated with Lys-C at a 1:50 (w/w) ratio (enzyme/protein) for 5 h, and then trypsin was added at a 1:50 (w/w) ratio (enzyme/protein) and the mixture was incubated overnight. The reaction was quenched by adding 20% TFA to a final concentration of ∼1%. Peptides were desalted on the AssayMAP Bravo platform using the peptide cleanup v2.0 protocol. C18 cartridges (Agilent, 5 µL bed volume) were primed with 100 µL 90% acetonitrile (ACN) and equilibrated with 70 µL 0.1% TFA at a flow rate of 10 µL/min. The samples were loaded at 5 µL/min, followed by an internal cartridge wash with 0.1% TFA at a flow rate of 10 µL/min. Peptides were eluted with 30 µL 80% ACN, 0.1% TFA, and dried in a Speed-Vac (Eppendorf) prior to TMT labeling.

For the phosphoproteomic analysis, protein digestion and C18 peptide cleanup were repeated on the protein lysates. After Speed-Vac, the peptides were resuspended in 80% ACN, and 0.1% TFA prior to the phosphopeptide enrichment.

#### TMT 11 plex labeling

The peptide amount in each sample was estimated using a quantitative colorimetric peptide assay kit (Thermo Fisher Scientific). Within each batch, equal amounts of peptides were labelled with TMT11 plex reagents, using a ratio of 0.8 mg reagent to 100 μg peptides. The TMT labeling was performed according to the manufacturer’s instructions. Peptides were resuspended in 100 µL of 200 mM TEAB and individual TMT11 plex reagents were dissolved in 41 µL of anhydrous ACN and mixed with the peptide solution. The internal reference sample, a pool consisting of aliquots from protein lysates from 40 melanoma patient samples, was labeled in channel 126 in each batch. After one hour of incubation, the reaction was quenched by adding 8 µL of 5% hydroxylamine and incubated at room temperature for 15 minutes. The labeled peptides were mixed in a single tube, the volume was reduced in a Speed-Vac and then the peptides were cleaned up using a Sep-Pak C18 96-well Plate (Waters). The eluted peptides were dried in a Speed-Vac and finally resuspended in water prior to high pH RP-HPLC fractionation. The samples were distributed among 15 batches, using TMT tag 126 as the internal reference sample as described in Table S1C.

#### High pH RP-HPLC fractionation

The TMT11 batches were fractionated using an Aeris Widepore XB-C8 (3.6 μm, 2.1 × 100 mm) column (Phenomenex) on an 1100 Series HPLC (Agilent) operating at 80 µL/min. The mobile phases were solvent A: 20 mM ammonium formate pH 10, and solvent B: 80% ACN and 20% water containing 20 mM ammonium formate pH 10. An estimated amount of 200 µg was separated using the following gradient: 0 min 5% B; 1 min 20% B; 60 min 40% B; 90 min 90% B; 120 min 90% B. The column was operated at RT and the detection wavelength was 220 nm. Then, 96 fractions were collected at 1 min intervals and further concatenated to 24 or 25 fractions (by combining 4 fractions that were 24 fractions apart so that #1, #25, #49, and #73; and so forth, were concatenated), and dried in a Speed-Vac.

#### Automated phosphopeptide enrichment

The Phospho Enrichment v2.0 protocol on the AssayMAP Bravo platform ^123^ was used to enrich phosphorylated peptides using 5 µL Fe(III)-NTA cartridges. The cartridges were primed with 100 µL 50% ACN, 0.1% TFA at a flow rate of 300 µL/min and equilibrated with 50 µL loading buffer (80% ACN, 0.1% TFA) at 10 µL/min. Samples were loaded onto the cartridge at 3.5 µL/min. The samples were washed with 50 µL loading buffer and the phosphorylated peptides were eluted with 25 µL 5% NaOH directly into 10 µL 50% formic acid. Samples were dried in a Speed-Vac and stored at -80°C until analysis by LC-MS/MS.

### MS data acquisition

#### nLC-MS/MS analysis

The nLC-MS/MS analysis was performed on an Ultimate 3000 HPLC coupled to a Q Exactive HF-X mass spectrometer (Thermo Scientific). Each fraction (1 µg) was loaded onto a trap column (Acclaim1 PepMap 100 pre-column, 75 µm, 2 cm, C18, 3 mm, 100 Å, Thermo Scientific) and then separated on an analytical column (EASY-Spray column, 25 cm, 75 µm i.d., PepMap RSLC C18, 2 mm, 100Å, Thermo Scientific) using solvent A: 0.1% formic acid in water and solvent B: 0.1% formic acid in ACN, at a flow rate of 300 nL/min and a column temperature of 45°C. An estimated peptide amount of 1 µg was injected into the column and the following gradient was used: 0 min 4% B; 3 min 4% B; 109 min 30% B; 124 min 45% B; 125 min 98% B; 130 min 98% B. The TMT node was utilized as follows: full MS scans at m/z 350-1,400 with a resolution of 120,000 at m/z 200, a target AGC value of 3×10^6^ and IT of 50 ms, DDA selection of the 20 most intense ions for fragmentation in HCD collision cell with an NCE of 34 and MS/MS spectra acquisition in the Orbitrap analyzer at a resolution of 45,000 (at m/z 200) with a maximum IT of 86 ms, fixed first mass of 110 m/z, isolation window of 0.7 Da and dynamic exclusion of 30 s.

#### Spectral library

Peptides were dissolved in 2% ACN, 0.1% TFA and spiked with iRT peptides (Biognosis AG) in a 1:10 dilution (iRT:peptides). First, a spectral library using DDA was built using the same LC-MS/MS system as the global proteome analysis with the same type of trap- and analytical column, flow rate, temperature, and solvents. The gradient used was the following: 0 min 4% B; 7 min 4% B; 139 min 30% B; 154 min 45% B; 155 min 98% B; 160 min 98% B. The MS parameters were set as follows: Selection of the 15 most intense ions for fragmentation, full MS scans at m/z 375-1,750 with a resolution of 120,000 at m/z 200, a target AGC value of 3×10^6^ and IT of 100 ms, fragmentation in HCD collision cell with normalized collision energy (NCE) of 25 and MS/MS spectra acquisition in the Orbitrap analyzer at a resolution of 60,000 (at m/z 200) with a maximum IT of 120 ms and dynamic exclusion of 30 s.

#### Phospho-DIA

For DIA-MS, the phosphopeptides were separated using the same gradient and MS system as for the DDA analysis and the iRT mix was added to the individual samples. The full scans were processed in the Orbitrap analyzer with a resolution of 120,000 at (200 m/z), an injection time of 50 ms, and a target AGC value of 3×10^6^ in a range of 350 to 1,410 m/z. Fragmentation was set to 54 variable isolation windows based on the density distribution of m/z precursors in the previously built spectral library. MS2 scans were acquired with a resolution of 30,000 at 200 m/z, an NCE of 25, a target AGC value of 1×10^6^, and 200 m/z as fixed first mass.

#### Quality control of the MS analysis

Quality control measurements were introduced to assess the performance of LC-MS/MS systems. A protein digest from HeLa cells (Pierce HeLa Protein Digest Standard, Thermo Fisher Scientific) mixed with a standard peptide mixture (Pierce Peptide Retention Time Calibration Mixture) was used as a QC sample and measured every tenth LC-MS/MS analysis. This allowed monitoring of the peak width, retention time, base peak intensity, number of MS/MS, Peptide-Spectrum matches (PSMs), and number of peptides and proteins identified.

### Immunohistochemistry

For the immunohistochemical study, 42 primary melanoma tissues were used. The fixation of the tumor material was performed after the surgical removal of the melanoma tissue with 4% cc. buffered formaldehyde (volume ratio 1 tissue/10 fixative). Samples were then dehydrated in xylene/ethanol solution and embedded in paraffin and stored at room temperature. Sections of 10 μm were used for further immunohistochemical analysis. Representative tissue areas from paraffin-embedded blocks were selected based on the HE-stained slides, then 5 mm circumferential columns were put into the tissue microarrays (TMAs) in an ordered manner. From TMAs, 3.5 µm sections were placed into an automated immunostainer (Leica Bond Max, Leica Biosystems) using standardized deparaffinization, rehydration, and staining processes based on the automatized “Polymer No Enhancer – Bond Polymer Refine IHC protocol no enhancer” protocol in Leica Bond Max (Leica Biosystems instrument).

Antibodies against ADAM10, CDK4, CTNND1, DDX11, FGA, HMOX1, SCAI, PAEP, and PIK3CS3 were applied in dilution series (Table S9B). For visualization, a high affinity polymer-based, AF-linked secondary antibody was used with a Fast Red chromogenic substrate. For negative controls, open containers were filled with primary antibody diluent without primary antibody. Before coverslipping, slides were counterstained with hematoxylin. All IHC stages were automatized.

The colorimetric immunostained slides were scanned by a 3D Histech slide scanner (Pannoramic MIDI, 2010 3D Histech Ltd.). The digitized images served as a basis for the densitometry quantification of the antibody expression using the Image Pro Plus software. Multicolor pictures were converted into a grayscale spectrum, and then representative areas of the cell cytoplasm and/or nucleus of both melanoma and stromal cells were measured separately based on the strongest antibody expression in the grayscale spectrum. The continuous scale variables were collected for statistical analysis.

### Proteomic data processing

#### TMT 11 plex quantification of proteomic data

The global proteomic experiment generated a total of 375 raw files that were processed with Proteome Discoverer 2.3 (Thermo Fisher Scientific) using the Sequest HT search engine. The search was performed against the Homo sapiens UniProt revised database (downloaded 2018-10-01) with isoforms. Cysteine carbamidomethylation (+57.0215 Da) and TMT6 plex (+229.1629 Da) at peptide N-terminus and lysine were set as fixed modifications while methionine oxidation (+15.9949 Da), N-terminal acetylation (+42.0105 Da) and were set as variable modifications; peptide mass tolerance for the precursor ions and MS/MS spectra were 10 ppm and 0.02 Da, respectively. A maximum of two missed cleavage sites was accepted and a maximum false discovery rate (FDR) of 1% was used for identification at PSM, peptide, and protein levels using all samples of the proteomic dataset. The Proteome Discoverer software allowed the introduction of reporter ion interferences for each batch of TMT11 plex reagents as isotope correction factors in the quantification method. The peptides that could be uniquely mapped to a protein were used for relative protein abundance calculations.

These search results were imported into the Perseus software v. 1.6.6.0 ^124^. To correct for experimental differences related to sample handling and other biases such as column changes, the protein intensities were log2 transformed and centered around zero by subtracting the median intensity in each sample. To allow for the comparison of relative protein abundances between the different batches of TMT11 plex the protein intensities from the pooled references sample (in channel 126 in each batch) were subtracted from each channel in the corresponding batch to obtain the final relative protein abundance values.

#### Evaluation of the TMT global proteomic data

The TMT-based proteomic analyses of 142 metastatic malignant melanoma samples resulted in the identification of 12,695 proteins (11,468 genes) with an average of 10,705 proteins identified per sample. The data displayed 15.5% missing values for the protein abundances and 8,124 proteins were present across all samples. Long-term reproducibility of the digestion workflow was previously shown ^125^. The reliability of the TMT workflow was evaluated by repeating the entire experiment of batch one. Although factors such as sample aging and change of RP-high pH fractionation column and MS instrument influenced the analysis, the overall agreement and correlation of protein abundances between the experiments were good (Figure S1C). In addition, good longitudinal performance across the 15 batches was demonstrated by the rather constant sequence coverage (Figure S1D).

Principal component analysis using 8,124 proteins quantified in all the 142 melanoma samples could separate between high-(>70%) and low-containing (<30%) tumor samples based on protein abundance (Figure S1E). No batch effects were observed for the global proteomic or phosphoproteomic data (Figures S1E and S1F). mRNA and protein abundances showed a strong positive correlation (median 0.408), and 84% showed a significant correlation (p<0.05) for the 6,101 overlapping genes across 104 patient samples (Figure S1G). The average correlation was in the range of previously reported mRNA-protein correlations from CPTAC proteogenomic studies.

### Phosphoproteomic data processing

#### DIA phosphopeptide quantification

The phosphoproteomic spectral library was generated from 45 DDA raw files in the Spectronaut X platform (Biognosis AG) ^126^ against the Homo sapiens database from Uniprot (downloaded 2019-01-15) with isoforms. The following parameters were used: cysteine carbamidomethylation (+57.0215 Da) as fixed modification and methionine oxidation (+15.9949 Da), N-terminal acetylation (+42.0105 Da) and phosphorylation (+79.9663 Da) on serine, threonine, and tyrosine were selected as variable modifications. A maximum of two missed cleavages were accepted. Precursor mass tolerance was set to 10 ppm and for the MS/MS fragments, it was set to 0.02 Da. Between 3 and 25 fragments were collected per peptide. The phosphosite localization algorithm was set so that phosphosites with a score equal to or higher than 0.75 (i.e., 75% accuracy of determination of the phosphate localization) were considered as Class I. Filtering was performed at 1% FDR at PSM, peptide and protein levels for the whole phosphoproteomic dataset.

The 122 DIA raw files were analyzed in Spectronaut X. In the transition settings, charges +2 and +3 were set for the precursor ions, and +1, +2, and +3 were set for the b- and y-ion products, with a mass tolerance of 0.02 Da. Both the precursor and protein q-value cutoffs were set to 0.01 and the peptides were quantified based on the intensity of the MS1 signal precursor. In all samples, the retention time alignment was performed with spiked-in iRT peptides (Biognosys). From the database search, a total of 45,356 phosphosites in 29,484 phosphopeptides were identified with an average of 18,722 phosphosites identified per sample. The data displayed 58.7% missing values in the phosphosite abundance. The data were exported into the Perseus software v. 1.6.2.3 ^124^. Valid value filtering was applied and all phosphosites with more than 5% missing values were removed. The data were then log2-transformed and centered around zero by subtracting the median intensity in each sample. For those phosphosites with less than 5% missing values, the phosphosite abundance values were imputed by applying the K Nearest Neighbor method, resulting in 4,644 phosphosites in each patient used for the kinase analysis, ICA, and survival analyses.

#### Evaluation of the phosphoproteomic data

The sample preparation workflow was previously assessed for its reliability using malignant melanoma tissue samples ^123^. Principal component analysis, using 1,267 phosphosites commonly quantified among 118 patient samples showed similar separation as for the global proteomic dataset, separating the high-(>70%) and low-containing (<30%) tumor samples based on phosphosite abundance (Figure S1F). Protein and phosphoprotein abundances showed a strong positive correlation (median 0.506), and a 94% significant correlation (p<0.05) for the 809 overlapping proteins across 94 patient samples (Figure S1H).

### Statistical analysis of omics data

#### Sample exclusion

Subtypes: >30% tumor content, n=118

Analysis of phenotype switching markers: >70% tumor content, n=83

ICA: >30% tumor content

BRAF V600: n = 49

TME: <50% tumor content, n= 29

Survival analyses (Outlier and Cox): >30% tumor content

Sample MM-SEG-0113 was excluded from all analyses due to technical issues.

#### Consensus clustering analyses

Unsupervised-consensus hierarchical clustering analysis was performed using the 3,000 proteins with the most variable expression levels (coefficient of variation > 0.36) using the Perseus software (v 1.6.14.0) ^124^. The clustering algorithm used k-means, Pearson correlation distance, and average linkage. Five subgroups were identified as proteomic subtypes by visual inspection of the hierarchical tree.

#### Subtypes and feature correlation

Correlations between subtypes and clinico-histopathological features and transcriptomic subtypes were performed using Fisher’s exact test. Kruskal-Wallis tests were performed on the histological parameters including the contents of tumor cells, adjacent lymph nodes, tumor-associated connective tissue, and necrosis.

#### Differential omics analysis

To identify proteins, phosphosites, and transcripts differentially expressed across the subtypes, one-way ANOVA was performed using the TMT-based global proteomic, phosphoproteomic, and transcriptomic data, respectively. At least 30% of valid values were required for all datasets and the p-values were adjusted using the permutation-based FDR method.

#### Independent component analysis

Pre-processed and normalized proteomic, transcriptomic, and phosphoproteomic data were dimensionally reduced by independent component analysis (ICA) separately 127. To ensure the quality of the ICA, we only included omics data of samples with tumor content above 30%. An R-based package, “fastICA”, was used for implementation. The ICA was performed 100 times for each omics dataset to make sure that the ICs were consistent. The extracted independent components (ICs) mixing scores of the omics data were then passed through association tests with the joint table of clinical features of patients in our cohort. If the clinical variable is binary, a logistic regression model was built for association tests. Otherwise, a linear regression model was built. The association tests were conducted for all the 100 ICA analyses for each omics dataset and its ICs, and the ICs showing correlations (p-value<0.005) with a clinical feature for at least 30 ICA runs per 100 were picked as significant. For each of these significant ICs, the centroid of IC coefficients was used to rank the omics data. We then used these rankings to conduct Gene Set Enrichment Analysis (GSEA) and significant pathways were found (adjusted p-value<0.01). The GSEA was implemented by an R-based package, “fgsea”, and searched against the “Reactome” database. The ICs served as links between clinical and histological features and pathways.

#### Identification of mortality risk subgroups of BRAF V600 mutated patients

The R package ‘InGRiD’ (Integrative Genomics Robust iDentification of cancer subgroups)^130^ was utilized to identify subgroups of patients with different mortality risk rates within a cohort of 49 patients with a BRAF mutation (Table S1A). The package provides a pathway-guided identification of patient subgroups based on protein expression while utilizing patient survival information as the outcome variable. The analysis was supplied with the relative abundances of proteins differentially expressed between tumors with high and low expression of the mutation, and the associated pathways. These proteins were extracted from a previous study ^131^. As the outcome variable, we considered the patient survival time from sample collection to death or censoring (DSS). All default parameters in ‘InGRiD’ were kept.

#### Custom database construction and Single Amino Acid Variant (SAAV) peptide identification

A custom protein sequence database was built by downloading protein mutation data from the Cancer Mutant Proteome Database ^132^. This included the skin cutaneous melanoma data from TCGA (369 cases) and 7 melanoma cell lines from the NCI-60 panel. Additional data on melanoma was retrieved from COSMIC v80 (downloaded 2017-02-15). Protein IDs, mutation or variant positions, and mutated protein sequences were extracted, and a UniProt ID was assigned to the proteins, using a custom script written in Tcl. Using the matched protein sequence, a peptide that carried the mutation site was generated by performing an in silico tryptic digestion of the protein and allowing for one additional missed cleavage at both sides of the mutation site. Redundant mutations were then removed and entries with the same mutated peptide sequence were grouped into one single entry. The resulting database comprised 57,134 entries.

Raw files were processed with Proteome Discoverer 2.3 using the Sequest HT search engine in a two-step search. The first search was performed against the *Homo Sapiens* Swissprot database (see “TMT11 plex quantification of proteomic data”), and unassigned MS/MS spectra were searched against the above-described in-house built database. Cysteine carbamidomethylation was set as fixed modification while methionine oxidation and TMT11 plex at peptide N-terminus and lysine were set as variable modifications; peptide mass tolerance for the precursor ions and MS/MS spectra was set to 10 ppm and 0.02 Da, respectively. A maximum of two missed cleavage sites were accepted and FDR was set at 1% for identification at the peptide level.

#### Validation of SAAV search results

SAAV peptides were validated using the SpectrumAI quality control tool available as an R script ^133^. A custom R script was used for data cleanup and post-processing. The verified SAAVs pointing at the same mutation position on a protein were merged into one entry. The reason for multiple entries includes missed cleavages as well as complementary peptides pointing at the same mutation. The latter occurred if the amino acid change generated a new trypsin cleavage site leading to a peptide that cannot be predicted from the original canonical sequence.

For peptides assigned to an isoform of the master protein, the mutation positions were corrected to reflect the position in the canonical Uniprot sequence. This was performed by using a customized R script analyzing the UniProtKB isoform sequences (accessed 2019-08-21). Both the corrected and uncorrected position was used for online database searches, to ensure that we would not miss matching results due to the position disparity caused by isoform sequences. Additionally, the Peptide-Spectrum Matches (PSMs) matching wild-type peptides originating from the normal database search were linked to the corresponding SAAVs, which allowed assessing the ratio of wild-type and SAAV peptide PSMs.

KEGG and GO enrichment analyses were done using the clusterProfiler R package ^134^. Signaling pathway members were obtained from KEGG ^135^, and UniProt accession IDs were converted to KEGG IDs using the KEGG Mapper ^136^ ^137–139^.

#### Annotation of validated SAAVs

Merging the results of various database searches and cleaning the data was performed with in-house custom R scripts. First, the SAAV peptides were searched in PeptideAtlas database ^140^ to find out if the SAAV peptides were observed previously. The search was performed on the webpage https://db.systemsbiology.net/sbeams/cgi/PeptideAtlas/GetPeptides using the “Human 2020-01” Atlas Build, only keeping the canonical and isoform protein accession numbers for which SAAVs were identified in our study. The resulting peptide sequences were downloaded in text format and a custom R script was used to retrieve exact and partial matches.

Validated coding SNPs and cancer-related mutations were downloaded from the CanProVar database ^141,142^. The UniProt IDs were first converted to Ensembl IDs using the biomaRt (version 2.42.1) R package ^143,144^, and then the CanProVar database was used to retrieve the variant’s reference SNP ID (rs#) and any cancer-related variation ID of CanProVar. The “Index of human polymorphisms and disease mutation” document was downloaded from UniProt (https://www.uniprot.org/docs/humsavar) and was also used to retrieve reference SNP IDs.

Validated SAAVs and the corresponding genes were searched for in The Cancer Gene Census (CGC) ^145,146^. Additionally, we used the bioinformatic predictor FATHHM (functional analysis through hidden Markov models, http://fathmm.biocompute.org.uk/cancer.html, Hum. Mutat., 34:57-65) using the recommended Prediction Threshold of -0.75 to look for cancer-associated variants in our SAAV dataset.

#### Signature of melanoma-associated SAAVs

Due to the lack of genomic and mutational data on individual tumor samples, a signature of 167 melanoma-associated SAAVs was extracted. The signature was classified into levels 1-3 and included SAAVs with markedly different occurrences in the melanoma samples compared to the healthy tissue of patients.

Data on aggregated allele frequency, named here as Variant Allele frequency (VAf) were accessed using the NCBI Variation Service API as described in https://github.com/ncbi/dbsnp/blob/master/tutorials/Variation%20Services/Jupyter_Notebook/by_rsid.ipynb ^147^. For this analysis a custom Python script was used. Additional resources such as ExAc ^148^, 1000Genomes ^149^, and HapMap ^150^ were used to manually retrieve VAf information when this information was missing. VAf in the European population (AAFs) were manually imputed for BRAF V600E, NRAS Q61K/R, HRAS G13D and CDKN2A P114L. For BRAF V600E, HRAS G13D and CDKN2A P114L the global population alternative allele frequency was used. VAfs for NRAS mutations were inferred both from BRAF V600E frequency and melanoma COSMIC data. According to COSMIC, BRAF mutation occurs 2.6471 times more than NRAS mutation (45% and 17% of the melanoma patients has these gene mutations, respectively), as well as NRAS Q61R occurs 1.067 times more frequently than NRAS Q61K (as 784 and 735 patients had Q61R and Q61K mutations, respectively). Based on these ratios, alternative allele frequency in the European population was inferred as AAF_BRAF Q61R_ = 3.1E-06 and AAF_NRAS Q61K_ = 2.9E-06.

#### Level 1

First, we estimated the frequency of SAAV (SAAVf) in the tumors as:

SAAVf = (n_SAAV PSM_) / (n_SAAV PSM_ + n_wild-type PSM_), where n_SAAV PSM_ is the number of PSMs supporting the SAAV peptide, while n_wild-type PSM_ is the number of PSMs supporting the wild-type peptide (Table S6A). The SAAVf displayed a significant correlation (Spearman 0.75, p-value=2.2e-16) with the corresponding Vaf (Figure S4E). Next, an enrichment factor SAAVr was defined as: SAAVr = SAAVf/ VAfE, where VAfE is the alternative allele frequency in the European population. Log_10_-transformed SAAVr values were subjected to Johnson transformation using Minitab (vs 17) to achieve values following the normal distribution (*M* = 0.01304, *SD* = 0.9988). The significance level was set to *α* = 0.1, and values outside of the range [-1.630; 1.656] were considered under-(n=49) or over-represented (n=34) SAAVs (Figure 5B and Table S6A).

#### Level 2

In total, 22 SAAVs (Table S6A) defined as: 1) VAfE < 0.3; 2) no detection of the wild-type peptide in our TMT experiment; and 3) evidence of the wt peptides in the Peptide Atlas database (Figure 5C and Table S6A). Thus, they likely constitute over-represented SAAVs in melanoma tumors.

#### Level 3

In total, 62 SAAVs (Table S6A) for which VAf was not available and could not be estimated (Figure 5C and Table S6A). In eight of the cases, the wt-peptide was not detected despite being identified in proteomic studies reported in the Peptide Atlas database. The absence of VAf data could be seen as an indicator of low occurrence in the population, with a potential association with melanoma.

#### Discovery of melanoma SAAV-based neoantigens

Peptides-containing SAAVs and their position within respective protein sequences were aligned with a database of a melanoma-associated immunopeptidome ^85^. The NetMHCpan-4.1 tool ^86^ was used to evaluate the MHC I binding of database-matched peptides and SAAV-altered peptide counterparts.

#### Tumor microenvironment analyses

The adjacent lymph node (LN) and tumor-derived connective tissue (CT) contents separated the cohort into two groups each: the high LN group (HLN) with LN > 27% and the low LN group (LLN) with LN ≤ 27%, and the high CT (HCT) group with CT > 45.5%, and the low CT group (LCT) with CT ≤ 45.5%. Connective tissue includes tumor-derived stroma and adipose tissue. The cut-off values were selected by ROC curves based on the ability of these histological features to discriminate between long and short survivals, considering a three-year survival from sample collection (DSS), and below 50% tumor content. Kaplan-Meier survival analysis with log-rank (Mantel-Cox) and Gehan-Breslow-Wilcoxon testing was used for univariate analysis between these groups. P-value < 0.05 was considered statistically significant, and values for patients reported as alive, with an “unknown cause of death” or “dead due to other reasons” were censored. These analyses were performed using Graph Pad Prism version 9.1.1.

#### Survival biomarker analyses

Two complementary supervised approaches were used to relate omics data to survival. First, Outlier analysis, which treats survival as a binary variable. Second, Cox analysis, considers survival as a continuous variable.

#### Outlier Analysis

Outlier analyses were performed for different variables of interest including survival, BRAF mutation, NRAS mutation, gender, and tumor stage. For the survival-related variables, the dataset was divided into 2 groups, and binary variables were created based on whether the patients lived longer than certain cutoff times, or not. The cutoff times used were 6 months, 1 year, 3 years, and 5 years from their sample collection date (DSS). Outlier analyses were conducted to find which proteins were significantly enriched in one group. We used a Python-based package, “BlackSheep”, to implement these analyses on the proteomic, transcriptomic, and phosphoproteomic data separately with a default median and interquartile range (IQR) of 1.5 ^151^. Significant genes were picked by an FDR cutoff at 5% in the group-wise comparisons. Proteins labeled as outliers in less than 30% of the patient samples in one group were excluded from the group-wise comparisons.

#### Cox’s proportional hazards survival analysis

In addition, we performed survival analysis using regularized Cox regression in a similar manner as previously published ^152^. The samples were randomly split into a training and a test set (80-20 training-test set). Using the training samples, a univariate Cox model was fitted for each feature individually and the 30 features with the lowest univariate p-values were selected and used as input to an elastic-net Cox model (Table S8B). The C-index was computed on the left-out test samples. This procedure was repeated 100 times for each omics dataset. We then considered the features that were selected by the Cox model in at least 50 of the 100 repetitions as significant and investigated these further ^152^.

A C-index of 1.0 means that the model “ranks” the samples perfectly, i.e., patients with a higher risk score (hazard score) died earlier than those with lower scores. A C-index of 0.5 is the expected performance of a random model. The C-index parameter is analogous to the AUC parameter (Area Under Curve) used for a binary classifier. The predictive power of our molecular data was moderate, and concordance indices (C-indices) varied between 0.538 and 0.601.

#### ROC curve analysis

The survival data of the patients were divided at 6 months, 1 year, 3 years, and 5 years into binary variables. Univariate receiver operating characteristic (ROC) curves of each binary survival variable and each protein expression were constructed by the ‘pROC’ package in R. Area under the ROC curve (AUC) was used as a measurement to determine the correlation between survival and the expression of specific proteins. For each protein, the cutoff point of expression that gave the maximum sum of sensitivity and specificity was used to divide the samples into a high-expression group and a low-expression group. Using the R package called ‘survival’, Kaplan-Meier curves were then introduced to reveal the survival differences between samples in these groups. The Kaplan-Meier p-values (log-rank test) were also calculated with ‘survminer’ in R, which was used as another statistical value to evaluate the relationship between survival and the expression of specific proteins.

#### Literature background of the biomarkers

Of the three proteins highlighted by specific phosphosites, several links to melanoma were found in the literature. ADAM10, a member of a family of endopeptidases with broad specificity, is involved in the membrane shedding of several proteins. Interestingly, ADAM10 together with ADAM17 may promote membrane shedding of immunosuppressive proteins such as PDL1 and LAG3 ^110–112^. FGA, is a part of the glycoprotein fibrinogen which has major functions in hemostasis, wound healing and also in immune responses. High plasma levels of fibrinogen have been attributed to poor prognosis in lung cancer and also in melanoma. PTM state of fibrinogen has been linked to the disease state of cancer ^153–160^. The third selected protein, HMOX1, is an antioxidant and anti-inflammatory enzyme involved in generating biliverdin and bilirubin. HMOX1 may promote cancer cell growth, tumor cell survival and resistance to treatment. HMOX1 has earlier been linked to a poor outcome of melanoma ^161–164^.

Six other proteins were selected for IHC analysis based on regulation at the protein level: SCAI is a highly conserved protein that acts on the RhoA–Dia1 pathway to regulate invasive cell migration. SCAI is downregulated in many human tumors and high expression of SCAI correlates with better survival in patients with breast and lung cancers ^114,115^; there is so far little information about SCAI in melanoma.

CDK4 is a well-known cancer target that regulates cell cycle and proliferation. In melanoma, mutations and dysregulation are commonly seen in CDK4 and proteins in its pathways, and several candidate drugs are in the clinical phase ^104–106^.

CTNND1 is a key regulator of cell-cell adhesion. Several studies suggest a link to melanoma. The long isoform (1A) was found regulated in our study. Longer isoforms, often enriched in tumors, have been reported as pro-tumorigenic, playing a role in EMT ^165–170^.

The helicase DDX11 has a role in chromatid cohesion. DDX11 has been reported upregulated with progression from noninvasive to invasive melanoma, and expressed at high levels in advanced melanoma ^116–119^.

Glycodelin or PAEP, is a secreted glycoprotein that regulates critical steps during fertilization and has immunomodulatory effects. PAEP is expressed in melanoma and involved in tumor proliferation, migration and may promote development of immune tolerance in tumors. PAEP expression is regulated in part by MITF ^171–175^.

PIK3CB, is part of the PI3K–AKT cascade which is one of the most studied pathways in cancer. This pathway has a role in cell survival, migration and in oncogenic transformation. In melanoma, PI3K–AKT pathway may be activated by mutations in NRAS or loss of suppressor PTEN ^176–178^.

#### Paired correlation analyses

To study the protein-based correlation between the primary melanoma and the metastasis, IHC analysis of the markers was also performed in nine additional metastases (Table S1B). First, a one-sample Kolmogorov-Smirnow test was conducted to determine the normality of the antibody expression data. For the comparison, Wilcoxon signed-rank test was used to examine whether proteins were differentially expressed between the tumors and the matched metastases. P-values of One-Sample Kolmogorov-Smirnow test and Wilcoxon signed-rank test were calculated by the IBM SPSS statistics package (26.0 version) software. P < 0.05 was considered statistically significant.

#### Survival analysis

To show the predictive impact of the nine markers, we performed independent t-tests to assess the differences in the means of the protein IHC expression values, quantified in the tumor cell and stroma parts, between the tumors of patients with OS of more than five years and less than five years. The assumption of homogeneity of variances was tested by Levene’s Test of Equality of Variances. For the cases where significant differential expression was obtained, univariate receiver operating characteristic (ROC) curve analysis was used to generate cutoff points for each protein that separated groups of patients with differences in OS. This was followed by Kaplan-Meier survival analyses based on models generated by the optimal cutpoint of each protein (Table S9A).

Independent t-test, ROC curve, Kaplan-Meier survival analysis, and figures including box plots showing p-values, quartile values, mean values, and 95% confidence intervals were produced by GraphPad IBM SPSS statistics package (version 26.0). P < 0.05 was considered statistically significant. Only samples from patients with OS > or <5 years (n=34) were included in the Mann-Whitney U t-test and ROC curve analyses.

## DATA AND CODE AVAILABILITY

- MS global proteomic data and phosphoproteomic (PXD041340) data have been deposited and are publicly available as of the date of publication.
- Transcriptomic data are publicly available https://doi.org:10.18632/oncotarget.3655
- Histology images reported in this paper will be shared by the lead contact upon request.
- All original code has been deposited to the Github repository: https://github.com/rhong3/Segundo_Melanoma and https://github.com/bszeitz/MM_Segundo and is publicly available as of the date of publication.
- Any additional information required to reanalyze the data reported in this paper is available from the lead contact upon request.

## ACKNOWLEDGMENTS

This study was supported by grants from the Berta Kamprad Foundation, and the Mats and Stefan Paulsson Trust, Lund, Sweden. This work was done under the auspices of a Memorandum of Understanding between the European Cancer Moonshot Center in Lund and the U.S. National Cancer Institute’s International Cancer Proteogenome Consortium (ICPC). ICPC encourages international cooperation among institutions and nations in proteogenomic cancer research in which proteogenomic datasets are made available to the public. This work was also done in collaboration with the U.S. National Cancer Institute’s Clinical Proteomic Tumor Analysis Consortium (CPTAC). Further support came from the National Cancer Institute (NCI) CPTAC grants U24CA210972 to D.F. L.V.K. is a recipient of the János Bolyai Research Scholarship of the Hungarian Academy of Sciences and is supported by the the Hungarian National Research, Development and Innovation Office (OTKA FK138696) and Semmelweis University STIA-KFI2021 grants. I.N.B. was supported by the Hungarian Academy of Sciences, the grant of OTKA-125509. L.S. was supported by the ÚNKP-21-3-SZTE-102 New National Excellence Program of the Ministry for Innovation and Technology from the source of the National Research, Development and Innovation Fund (University of Szeged, Szeged, Hungary). B.S. was supported by the Semmelweis 250+ Excellence Ph.D. Scholarship (EFOP-3.6.3-VEKOP-16-2017-00009) and the ÚNKP-22-3-II New National Excellence Program of the Ministry for Culture and Innovation from the source of the National Research, Development and Innovation Fund. The authors are sincerely grateful to Professor Håkan Olsson (SUS University Hospital Lund) who was involved in the initiation of the study but unfortunately died prior to the submission of this manuscript. We also thank Nicole Woldmar and Felipe Sieira for designing the graphical abstract.

## AUTHOR CONTRIBUTIONS

The project was conceived and supervised by L.H.B., I.B.N., D.F, G.M. Clinical data review and inclusion of patients was conducted by I.B.N., H.E., B.B., C.I., H.O., L.L., H.L., C.W., and M.R. Sample preparation and generation of proteomic and phosphoproteomic data was performed by M.K., J.E., L.H.B. Pathological evaluation and immunohistochemistry was performed by A.M.S., L.S., I.B.N. Statistical analyses and biological interpretation was conducted by M.K., L.H.B., R.H., B.S., L.S., B.D., P.H., I.P., J.E., A.S., Z.H., J.G., L.V.K., M.R., K.P., E.W., D.F. The original draft was written by M.K., L.H.B., R.H., L.S., P.H., A.M.S., K.P., E.W., D.F., I.N. and G.M. Project administration was performed by R.A, J.M, and G.M. All authors edited or commented on the manuscript.

## REFERENCES

1. Thomas, D., and Bello, D.M. (2021). Adjuvant immunotherapy for melanoma. J. Surg. Oncol. 123, 789–797.

2. Boussios, S., Rassy, E., Samartzis, E., Moschetta, M., Sheriff, M., Pérez-Fidalgo, J.A., and Pavlidis, N. (2021). Melanoma of unknown primary: New perspectives for an old story. Crit. Rev. Oncol. Hematol. 158, 103208.

3. Bertolotto, C. (2013). Melanoma: from melanocyte to genetic alterations and clinical options. Scientifica 2013, 635203.

4. Grzywa, T.M., Paskal, W., and Włodarski, P.K. (2017). Intratumor and Intertumor Heterogeneity in Melanoma. Transl. Oncol. 10, 956–975.

5. Willmore-Payne, C., Holden, J.A., Hirschowitz, S., and Layfield, L.J. (2006). BRAF and c-kit gene copy number in mutation-positive malignant melanoma. Hum. Pathol. 37, 520–527.

6. Kiuru, M., and Busam, K.J. (2017). The NF1 gene in tumor syndromes and melanoma. Lab. Invest. 97, 146–157.

7. Cancer Genome Atlas Network (2015). Genomic Classification of Cutaneous Melanoma. Cell 161, 1681–1696.

8. Lo, J.A., and Fisher, D.E. (2014). The melanoma revolution: from UV carcinogenesis to a new era in therapeutics. Science 346, 945–949.

9. Ostrowski, S.M., and Fisher, D.E. (2021). Biology of Melanoma. Hematol. Oncol. Clin. North Am. 35, 29–56.

10. Kalbasi, A., and Ribas, A. (2020). Tumour-intrinsic resistance to immune checkpoint blockade. Nat. Rev. Immunol. 20, 25–39.

11. Darvin, P., Toor, S.M., Sasidharan Nair, V., and Elkord, E. (2018). Immune checkpoint inhibitors: recent progress and potential biomarkers. Exp. Mol. Med. 50, 1–11.

12. Cirenajwis, H., Ekedahl, H., Lauss, M., Harbst, K., Carneiro, A., Enoksson, J., Rosengren, F., Werner-Hartman, L., Törngren, T., Kvist, A., et al. (2015). Molecular stratification of metastatic melanoma using gene expression profiling: Prediction of survival outcome and benefit from molecular targeted therapy. Oncotarget 6, 12297–12309.

13. Jönsson, G., Busch, C., Knappskog, S., Geisler, J., Miletic, H., Ringnér, M., Lillehaug, J.R., Borg, A., and Lønning, P.E. (2010). Gene expression profiling-based identification of molecular subtypes in stage IV melanomas with different clinical outcome. Clin. Cancer Res. 16, 3356– 3367.

14. Harel, M., Ortenberg, R., Varanasi, S.K., Mangalhara, K.C., Mardamshina, M., Markovits, E., Baruch, E.N., Tripple, V., Arama-Chayoth, M., Greenberg, E., et al. (2019). Proteomics of Melanoma Response to Immunotherapy Reveals Mitochondrial Dependence. Cell 179, 236–250.e18.

15. Betancourt, L.H., Gil, J., Sanchez, A., Doma, V., Kuras, M., Murillo, J.R., Velasquez, E., Çakır, U., Kim, Y., Sugihara, Y., et al. (2021). The Human Melanoma Proteome Atlas-Complementing the melanoma transcriptome. Clin. Transl. Med. 11, e451.

16. Betancourt, L.H., Gil, J., Kim, Y., Doma, V., Çakır, U., Sanchez, A., Murillo, J.R., Kuras, M., Parada, I.P., Sugihara, Y., et al. (2021). The human melanoma proteome atlas-Defining the molecular pathology. Clin. Transl. Med. 11, e473.

17. Betancourt, L.H., Pawłowski, K., Eriksson, J., Szasz, A.M., Mitra, S., Pla, I., Welinder, C., Ekedahl, H., Broberg, P., Appelqvist, R., et al. (2019). Improved survival prognostication of node-positive malignant melanoma patients utilizing shotgun proteomics guided by histopathological characterization and genomic data. Sci. Rep. 9, 5154.

18. Ajona, D., Ortiz-Espinosa, S., Pio, R., and Lecanda, F. (2019). Complement in Metastasis: A Comp in the Camp. Front. Immunol. 10, 669.

19. Mollinedo, F. (2019). Neutrophil Degranulation, Plasticity, and Cancer Metastasis. Trends Immunol. 40, 228–242.

20. Kobayashi, T., Ogawa, H., Kasahara, M., Shiozawa, Z., and Matsuzawa, T. (1995). A single amino acid substitution within the mature sequence of ornithine aminotransferase obstructs mitochondrial entry of the precursor. Am. J. Hum. Genet. 57, 284–291.

21. Maurya, S.R., and Mahalakshmi, R. (2017). Mitochondrial VDAC2 and cell homeostasis: highlighting hidden structural features and unique functionalities. Biol. Rev. Camb. Philos. Soc. 92, 1843–1858.

22. Wu, Z.-Z., Wang, S., Yang, Q.-C., Wang, X.-L., Yang, L.-L., Liu, B., and Sun, Z.-J. (2020). Increased Expression of SHMT2 Is Associated With Poor Prognosis and Advanced Pathological Grade in Oral Squamous Cell Carcinoma. Front. Oncol. 10, 588530.

23. Afshar-Kharghan, V. (2017). The role of the complement system in cancer. J. Clin. Invest. 127, 780–789.

24. Pio, R., Corrales, L., and Lambris, J.D. (2014). The role of complement in tumor growth. Adv. Exp. Med. Biol. 772, 229–262.

25. Rooney, M.S., Shukla, S.A., Wu, C.J., Getz, G., and Hacohen, N. (2015). Molecular and genetic properties of tumors associated with local immune cytolytic activity. Cell 160, 48–61.

26. Weinstein, D., Leininger, J., Hamby, C., and Safai, B. (2014). Diagnostic and prognostic biomarkers in melanoma. J. Clin. Aesthet. Dermatol. 7, 13–24.

27. Goding, C.R., and Arnheiter, H. (2019). MITF-the first 25 years. Genes Dev. 33, 983–1007.

28. Hoek, K.S., and Goding, C.R. (2010). Cancer stem cells versus phenotype-switching in melanoma. Pigment Cell Melanoma Res. 23, 746–759.

29. Fürst, K., Steder, M., Logotheti, S., Angerilli, A., Spitschak, A., Marquardt, S., Schumacher, T., Engelmann, D., Herchenröder, O., Rupp, R.A.W., et al. (2019). DNp73-induced degradation of tyrosinase links depigmentation with EMT-driven melanoma progression. Cancer Lett. 442, 299– 309.

30. Pedri, D., Karras, P., Landeloos, E., Marine, J.-C., and Rambow, F. (2021). Epithelial-to-mesenchymal-like transition events in melanoma. FEBS J. 10.1111/febs.16021.

31. Woods, K., Pasam, A., Jayachandran, A., Andrews, M.C., and Cebon, J. (2014). Effects of epithelial to mesenchymal transition on T cell targeting of melanoma cells. Front. Oncol. 4, 367.

32. Tirosh, I., Izar, B., Prakadan, S.M., Wadsworth, M.H., 2nd, Treacy, D., Trombetta, J.J., Rotem, A., Rodman, C., Lian, C., Murphy, G., et al. (2016). Dissecting the multicellular ecosystem of metastatic melanoma by single-cell RNA-seq. Science 352, 189–196.

33. Kohrman, A.Q., and Matus, D.Q. (2017). Divide or Conquer: Cell Cycle Regulation of Invasive Behavior. Trends Cell Biol. 27, 12–25.

34. Aisner, D.L., Maker, A., Rosenberg, S.A., and Berman, D.M. (2005). Loss of S100 antigenicity in metastatic melanoma. Hum. Pathol. 36, 1016–1019.

35. Blessing, K., Sanders, D.S., and Grant, J.J. (1998). Comparison of immunohistochemical staining of the novel antibody melan-A with S100 protein and HMB-45 in malignant melanoma and melanoma variants. Histopathology 32, 139–146.

36. Xiong, T.-F., Pan, F.-Q., and Li, D. (2019). Expression and clinical significance of S100 family genes in patients with melanoma. Melanoma Res. 29, 23–29.

37. Bresnick, A.R., Weber, D.J., and Zimmer, D.B. (2015). S100 proteins in cancer. Nat. Rev. Cancer 15, 96–109.

38. Chen, H., Xu, C., Jin, Q. ‘e, and Liu, Z. (2014). S100 protein family in human cancer. Am. J. Cancer Res. 4, 89–115.

39. Massi, D., Landriscina, M., Piscazzi, A., Cosci, E., Kirov, A., Paglierani, M., Di Serio, C., Mourmouras, V., Fumagalli, S., Biagioli, M., et al. (2010). S100A13 is a new angiogenic marker in human melanoma. Mod. Pathol. 23, 804–813.

40. Åberg, A.-M., Bergström, S.H., Thysell, E., Tjon-Kon-Fat, L.-A., Nilsson, J.A., Widmark, A., Thellenberg-Karlsson, C., Bergh, A., Wikström, P., and Lundholm, M. (2021). High Monocyte Count and Expression of S100A9 and S100A12 in Peripheral Blood Mononuclear Cells Are Associated with Poor Outcome in Patients with Metastatic Prostate Cancer. Cancers 13. 10.3390/cancers13102424.

41. Janka, E.A., Várvölgyi, T., Sipos, Z., Soós, A., Hegyi, P., Kiss, S., Dembrovszky, F., Csupor, D., Kéringer, P., Pécsi, D., et al. (2021). Predictive Performance of Serum S100B Versus LDH in Melanoma Patients: A Systematic Review and Meta-Analysis. Front. Oncol. 11, 772165.

42. Gassenmaier, M., Lenders, M.M., Forschner, A., Leiter, U., Weide, B., Garbe, C., Eigentler, T.K., and Wagner, N.B. (2021). Serum S100B and LDH at Baseline and During Therapy Predict the Outcome of Metastatic Melanoma Patients Treated with BRAF Inhibitors. Target. Oncol. 16, 197–205.

43. Donato, R., Cannon, B.R., Sorci, G., Riuzzi, F., Hsu, K., Weber, D.J., and Geczy, C.L. (2013). Functions of S100 proteins. Curr. Mol. Med. 13, 24–57.

44. Flockhart, D.A., and Corbin, J.D. (1982). Regulatory mechanisms in the control of protein kinases. CRC Crit. Rev. Biochem. 12, 133–186.

45. Smith, J.A., Francis, S.H., and Corbin, J.D. (1993). Autophosphorylation: a salient feature of protein kinases. Molecular and Cellular Biochemistry 127-128, 51–70. 10.1007/bf01076757.

46. Wang, Z.-X., and Jia-Wei, W.U. (2002). Autophosphorylation kinetics of protein kinases. Biochemical Journal 368, 947–952. 10.1042/bj20020557.

47. Krug, K., Jaehnig, E.J., Satpathy, S., Blumenberg, L., Karpova, A., Anurag, M., Miles, G., Mertins, P., Geffen, Y., Tang, L.C., et al. (2020). Proteogenomic Landscape of Breast Cancer Tumorigenesis and Targeted Therapy. Cell 183, 1436–1456.e31.

48. Hoeflich, K.P., Luo, J., Rubie, E.A., Tsao, M.S., Jin, O., and Woodgett, J.R. (2000). Requirement for glycogen synthase kinase-3beta in cell survival and NF-kappaB activation. Nature 406, 86–90.

49. Vashishtha, V., Jinghan, N., and K Yadav, A. (2018). Antagonistic role of GSK3 isoforms in glioma survival. J. Cancer 9, 1846–1855.

50. Madhunapantula, S.V., Sharma, A., Gowda, R., and Robertson, G.P. (2013). Identification of glycogen synthase kinase 3α as a therapeutic target in melanoma. Pigment Cell Melanoma Res. 26, 886–899.

51. El Masri, R., and Delon, J. (2021). RHO GTPases: from new partners to complex immune syndromes. Nat. Rev. Immunol. 21, 499–513.

52. Neurohr, G.E., Terry, R.L., Lengefeld, J., Bonney, M., Brittingham, G.P., Moretto, F., Miettinen, T.P., Vaites, L.P., Soares, L.M., Paulo, J.A., et al. (2019). Excessive Cell Growth Causes Cytoplasm Dilution And Contributes to Senescence. Cell 176, 1083–1097.e18.

53. Leikam, C., Hufnagel, A.L., Otto, C., Murphy, D.J., Mühling, B., Kneitz, S., Nanda, I., Schmid, M., Wagner, T.U., Haferkamp, S., et al. (2015). In vitro evidence for senescent multinucleated melanocytes as a source for tumor-initiating cells. Cell Death Dis. 6, e1711.

54. Litwiniec, A., Gackowska, L., Helmin-Basa, A., Zuryń, A., and Grzanka, A. (2013). Low-dose etoposide-treatment induces endoreplication and cell death accompanied by cytoskeletal alterations in A549 cells: Does the response involve senescence? The possible role of vimentin. Cancer Cell Int. 13, 9.

55. Buetow, K.H., Meador, L.R., Menon, H., Lu, Y.-K., Brill, J., Cui, H., Roe, D.J., DiCaudo, D.J., and Hastings, K.T. (2019). High GILT Expression and an Active and Intact MHC Class II Antigen Presentation Pathway Are Associated with Improved Survival in Melanoma. J. Immunol. 203, 2577–2587.

56. van Tuyn, J., Jaber-Hijazi, F., MacKenzie, D., Cole, J.J., Mann, E., Pawlikowski, J.S., Rai, T.S., Nelson, D.M., McBryan, T., Ivanov, A., et al. (2017). Oncogene-Expressing Senescent Melanocytes Up-Regulate MHC Class II, a Candidate Melanoma Suppressor Function. J. Invest. Dermatol. 137, 2197–2207.

57. Chen, H.-A., Ho, Y.-J., Mezzadra, R., Adrover, J.M., Smolkin, R., Zhu, C., Woess, K., Bernstein, N., Schmitt, G., Fong, L., et al. (2022). Senescence rewires microenvironment sensing to facilitate anti-tumor immunity. Cancer Discov. 10.1158/2159-8290.CD-22-0528.

58. Katlinskaya, Y.V., Katlinski, K.V., Yu, Q., Ortiz, A., Beiting, D.P., Brice, A., Davar, D., Sanders, C., Kirkwood, J.M., Rui, H., et al. (2016). Suppression of Type I Interferon Signaling Overcomes Oncogene-Induced Senescence and Mediates Melanoma Development and Progression. Cell Rep. 15, 171–180.

59. Raaijmakers, M.I.G., Widmer, D.S., Narechania, A., Eichhoff, O., Freiberger, S.N., Wenzina, J., Cheng, P.F., Mihic-Probst, D., Desalle, R., Dummer, R., et al. (2016). Co-existence of BRAF and NRAS driver mutations in the same melanoma cells results in heterogeneity of targeted therapy resistance. Oncotarget 7, 77163–77174.

60. Leon, K.E., and Aird, K.M. (2019). Jumonji C Demethylases in Cellular Senescence. Genes 10. 10.3390/genes10010033.

61. Ohta, K., Haraguchi, N., Kano, Y., Kagawa, Y., Konno, M., Nishikawa, S., Hamabe, A., Hasegawa, S., Ogawa, H., Fukusumi, T., et al. (2013). Depletion of JARID1B induces cellular senescence in human colorectal cancer. Int. J. Oncol. 42, 1212–1218.

62. Chicas, A., Kapoor, A., Wang, X., Aksoy, O., Evertts, A.G., Zhang, M.Q., Garcia, B.A., Bernstein, E., and Lowe, S.W. (2012). H3K4 demethylation by Jarid1a and Jarid1b contributes to retinoblastoma-mediated gene silencing during cellular senescence. Proc. Natl. Acad. Sci. U. S. A. 109, 8971–8976.

63. Bayo, J., Tran, T.A., Wang, L., Peña-Llopis, S., Das, A.K., and Martinez, E.D. (2018). Jumonji Inhibitors Overcome Radioresistance in Cancer through Changes in H3K4 Methylation at Double-Strand Breaks. Cell Rep. 25, 1040–1050.e5.

64. Li, X., Liu, L., Yang, S., Song, N., Zhou, X., Gao, J., Yu, N., Shan, L., Wang, Q., Liang, J., et al. (2014). Histone demethylase KDM5B is a key regulator of genome stability. Proc. Natl. Acad. Sci. U. S. A. 111, 7096–7101.

65. Xu, J., and Kidder, B.L. (2018). KDM5B decommissions the H3K4 methylation landscape of self-renewal genes during trophoblast stem cell differentiation. Biol. Open 7. 10.1242/bio.031245.

66. Mallette, F.A., and Richard, S. (2012). JMJD2A promotes cellular transformation by blocking cellular senescence through transcriptional repression of the tumor suppressor CHD5. Cell Rep. 2, 1233–1243.

67. Salomoni, P., and Pandolfi, P.P. (2002). The role of PML in tumor suppression. Cell 108, 165– 170.

68. Gresko, E., Ritterhoff, S., Sevilla-Perez, J., Roscic, A., Fröbius, K., Kotevic, I., Vichalkovski, A., Hess, D., Hemmings, B.A., and Schmitz, M.L. (2009). PML tumor suppressor is regulated by HIPK2-mediated phosphorylation in response to DNA damage. Oncogene 28, 698–708.

69. Lim, J.H., Liu, Y., Reineke, E., and Kao, H.-Y. (2011). Mitogen-activated Protein Kinase Extracellular Signal-regulated Kinase 2 Phosphorylates and Promotes Pin1 Protein-dependent Promyelocytic Leukemia Protein Turnover *. J. Biol. Chem. 286, 44403–44411.

70. Yuan, W.-C., Lee, Y.-R., Huang, S.-F., Lin, Y.-M., Chen, T.-Y., Chung, H.-C., Tsai, C.-H., Chen, H.-Y., Chiang, C.-T., Lai, C.-K., et al. (2011). A Cullin3-KLHL20 Ubiquitin ligase-dependent pathway targets PML to potentiate HIF-1 signaling and prostate cancer progression. Cancer Cell 20, 214–228.

71. Zhou, W., and Bao, S. (2014). PML-mediated signaling and its role in cancer stem cells. Oncogene 33, 1475–1484.

72. Shihab, H.A. fathmm - Analyze Cancer-Associated Variants. http://fathmm.biocompute.org.uk/cancer.html.

73. Shihab, H.A., Gough, J., Cooper, D.N., Stenson, P.D., Barker, G.L.A., Edwards, K.J., Day, I.N.M., and Gaunt, T.R. (2013). Predicting the functional, molecular, and phenotypic consequences of amino acid substitutions using hidden Markov models. Hum. Mutat. 34, 57–65.

74. Hayward, N.K., Wilmott, J.S., Waddell, N., Johansson, P.A., Field, M.A., Nones, K., Patch, A.-M., Kakavand, H., Alexandrov, L.B., Burke, H., et al. (2017). Whole-genome landscapes of major melanoma subtypes. Nature 545, 175–180.

75. Lin, J., Wang, J., Greisinger, A.J., Grossman, H.B., Forman, M.R., Dinney, C.P., Hawk, E.T., and Wu, X. (2010). Energy balance, the PI3K-AKT-mTOR pathway genes, and the risk of bladder cancer. Cancer Prev. Res. 3, 505–517.

76. Li, Q., Yang, J., Yu, Q., Wu, H., Liu, B., Xiong, H., Hu, G., Zhao, J., Yuan, X., and Liao, Z. (2013). Associations between single-nucleotide polymorphisms in the PI3K-PTEN-AKT-mTOR pathway and increased risk of brain metastasis in patients with non-small cell lung cancer. Clin. Cancer Res. 19, 6252–6260.

77. Qi, L., Sun, K., Zhuang, Y., Yang, J., and Chen, J. (2017). Study on the association between PI3K/AKT/mTOR signaling pathway gene polymorphism and susceptibility to gastric cancer. J. BUON 22, 1488–1493.

78. Izzi, V., Davis, M.N., and Naba, A. (2020). Pan-Cancer Analysis of the Genomic Alterations and Mutations of the Matrisome. Cancers 12. 10.3390/cancers12082046.

79. dbSNP Home Page http://www.ncbi.nlm.nih.gov/SNP.

80. Park, S., Supek, F., and Lehner, B. (2018). Systematic discovery of germline cancer predisposition genes through the identification of somatic second hits. Nat. Commun. 9, 2601.

81. Kronert, W.A., Bell, K.M., Viswanathan, M.C., Melkani, G.C., Trujillo, A.S., Huang, A., Melkani, A., Cammarato, A., Swank, D.M., and Bernstein, S.I. (2018). Prolonged cross-bridge binding triggers muscle dysfunction in a Drosophila model of myosin-based hypertrophic cardiomyopathy. Elife 7. 10.7554/eLife.38064.

82. Marston, S. (2018). The Molecular Mechanisms of Mutations in Actin and Myosin that Cause Inherited Myopathy. Int. J. Mol. Sci. 19. 10.3390/ijms19072020.

83. Orgaz, J.L., Crosas-Molist, E., Sadok, A., Perdrix-Rosell, A., Maiques, O., Rodriguez-Hernandez, I., Monger, J., Mele, S., Georgouli, M., Bridgeman, V., et al. (2020). Myosin II Reactivation and Cytoskeletal Remodeling as a Hallmark and a Vulnerability in Melanoma Therapy Resistance. Cancer Cell 37, 85–103.e9.

84. McGrail, D.J., Pilié, P.G., Rashid, N.U., Voorwerk, L., Slagter, M., Kok, M., Jonasch, E., Khasraw, M., Heimberger, A.B., Lim, B., et al. (2021). High tumor mutation burden fails to predict immune checkpoint blockade response across all cancer types. Ann. Oncol. 32, 661–672.

85. Bassani-Sternberg, M., Bräunlein, E., Klar, R., Engleitner, T., Sinitcyn, P., Audehm, S., Straub, M., Weber, J., Slotta-Huspenina, J., Specht, K., et al. (2016). Direct identification of clinically relevant neoepitopes presented on native human melanoma tissue by mass spectrometry. Nat. Commun. 7, 13404.

86. NetMHCpan - 4.1 NetMHCpan - 4.1. https://services.healthtech.dtu.dk/service.php?NetMHCpan-4.1.

87. Yuan, Y., Ju, Y.S., Kim, Y., Li, J., Wang, Y., Yoon, C.J., Yang, Y., Martincorena, I., Creighton, C.J., Weinstein, J.N., et al. (2020). Comprehensive molecular characterization of mitochondrial genomes in human cancers. Nat. Genet. 52, 342–352.

88. Venning, F.A., Wullkopf, L., and Erler, J.T. (2015). Targeting ECM Disrupts Cancer Progression. Front. Oncol. 5, 224.

89. Woo, H.Y., Rhee, H., Yoo, J.E., Kim, S.H., Choi, G.H., Kim, D.Y., Woo, H.G., Lee, H.S., and Park, Y.N. (2022). Lung and lymph node metastases from hepatocellular carcinoma: Comparison of pathological aspects. Liver Int. 42, 199–209.

90. Fitzgerald, A.A., and Weiner, L.M. (2020). The role of fibroblast activation protein in health and malignancy. Cancer Metastasis Rev. 39, 783–803.

91. Puré, E., and Blomberg, R. (2018). Pro-tumorigenic roles of fibroblast activation protein in cancer: back to the basics. Oncogene 37, 4343–4357.

92. Mesker, W.E., van Pelt, G.W., and Tollenaar, R.A.E.M. (2019). Tumor stroma as contributing factor in the lymph node metastases process? Oncotarget 10, 922–923.

93. van Pelt, G.W., Sandberg, T.P., Morreau, H., Gelderblom, H., van Krieken, J.H.J.M., Tollenaar, R.A.E.M., and Mesker, W.E. (2018). The tumour-stroma ratio in colon cancer: the biological role and its prognostic impact. Histopathology 73, 197–206.

94. Vangangelt, K.M.H., Tollenaar, L.S.A., van Pelt, G.W., de Kruijf, E.M., Dekker, T.J.A., Kuppen, P.J.K., Tollenaar, R.A.E.M., and Mesker, W.E. (2018). The prognostic value of tumor-stroma ratio in tumor-positive axillary lymph nodes of breast cancer patients. Int. J. Cancer 143, 3194–3200.

95. Röcken, M. (2010). Early tumor dissemination, but late metastasis: insights into tumor dormancy. J. Clin. Invest. 120, 1800–1803.

96. Buder-Bakhaya, K., and Hassel, J.C. (2018). Biomarkers for Clinical Benefit of Immune Checkpoint Inhibitor Treatment-A Review From the Melanoma Perspective and Beyond. Front. Immunol. 9, 1474.

97. Tsoi, J., Robert, L., Paraiso, K., Galvan, C., Sheu, K.M., Lay, J., Wong, D.J.L., Atefi, M., Shirazi, R., Wang, X., et al. (2018). Multi-stage Differentiation Defines Melanoma Subtypes with Differential Vulnerability to Drug-Induced Iron-Dependent Oxidative Stress. Cancer Cell 33, 890–904.e5.

98. Davis, L.E., Shalin, S.C., and Tackett, A.J. (2019). Current state of melanoma diagnosis and treatment. Cancer Biol. Ther. 20, 1366–1379.

99. Rambow, F., Marine, J.-C., and Goding, C.R. (2019). Melanoma plasticity and phenotypic diversity: therapeutic barriers and opportunities. Genes Dev. 33, 1295–1318.

100. Yuan, Y. (2016). Spatial Heterogeneity in the Tumor Microenvironment. Cold Spring Harb. Perspect. Med. 6. 10.1101/cshperspect.a026583.

101. Souza da Silva, R.M., Queiroga, E.M., Paz, A.R., Neves, F.F.P., Cunha, K.S., and Dias, E.P. (2021). Standardized Assessment of the Tumor-Stroma Ratio in Colorectal Cancer: Interobserver Validation and Reproducibility of a Potential Prognostic Factor. Clin Pathol 14, 2632010X21989686.

102. Hagenaars, S.C., Vangangelt, K.M.H., Van Pelt, G.W., Karancsi, Z., Tollenaar, R.A.E.M., Green, A.R., Rakha, E.A., Kulka, J., and Mesker, W.E. (2022). Standardization of the tumor-stroma ratio scoring method for breast cancer research. Breast Cancer Res. Treat. 193, 545–553.

103. Gao, J., Shen, Z., Deng, Z., and Mei, L. (2021). Impact of Tumor-Stroma Ratio on the Prognosis of Colorectal Cancer: A Systematic Review. Front. Oncol. 11, 738080.

104. Guo, L., Qi, J., Wang, H., Jiang, X., and Liu, Y. (2020). Getting under the skin: The role of CDK4/6 in melanomas. Eur. J. Med. Chem. 204, 112531.

105. Freedberg, D.E., Rigas, S.H., Russak, J., Gai, W., Kaplow, M., Osman, I., Turner, F., Randerson-Moor, J.A., Houghton, A., Busam, K., et al. (2008). Frequent p16-independent inactivation of p14ARF in human melanoma. J. Natl. Cancer Inst. 100, 784–795.

106. Hocker, T., and Tsao, H. (2007). Ultraviolet radiation and melanoma: a systematic review and analysis of reported sequence variants. Hum. Mutat. 28, 578–588.

107. Guan, X., LaPak, K.M., Hennessey, R.C., Yu, C.Y., Shakya, R., Zhang, J., and Burd, C.E. (2017). Stromal Senescence By Prolonged CDK4/6 Inhibition Potentiates Tumor Growth. Mol. Cancer Res. 15, 237–249.

108. Miller, M.A., Sullivan, R.J., and Lauffenburger, D.A. (2017). Molecular Pathways: Receptor Ectodomain Shedding in Treatment, Resistance, and Monitoring of Cancer. Clin. Cancer Res. 23, 623–629.

109. Smith, T.M., Jr, Tharakan, A., and Martin, R.K. (2020). Targeting ADAM10 in Cancer and Autoimmunity. Front. Immunol. 11, 499.

110. Andrews, L.P., Szymczak-Workman, A.L., Workman, C.J., and Vignali, D.A.A. (2015). The extent of metalloproteinase-mediated LAG3 cleavage limits the efficacy of PD1 blockade. J Immunother Cancer 3, P216.

111. Lambrecht, B.N., Vanderkerken, M., and Hammad, H. (2018). The emerging role of ADAM metalloproteinases in immunity. Nat. Rev. Immunol. 18, 745–758.

112. Orme, J.J., Jazieh, K.A., Xie, T., Harrington, S., Liu, X., Ball, M., Madden, B., Charlesworth, M.C., Azam, T.U., Lucien, F., et al. (2020). ADAM10 and ADAM17 cleave PD-L1 to mediate PD-(L)1 inhibitor resistance. Oncoimmunology 9, 1744980.

113. Andrews, L.P., Marciscano, A.E., Drake, C.G., and Vignali, D.A.A. (2017). LAG3 (CD223) as a cancer immunotherapy target. Immunol. Rev. 276, 80–96.

114. Brandt, D.T., Baarlink, C., Kitzing, T.M., Kremmer, E., Ivaska, J., Nollau, P., and Grosse, R. (2009). SCAI acts as a suppressor of cancer cell invasion through the transcriptional control of beta1-integrin. Nat. Cell Biol. 11, 557–568.

115. Gasparics, Á., Kökény, G., Fintha, A., Bencs, R., Mózes, M.M., Ágoston, E.I., Buday, A., Ivics, Z., Hamar, P., Győrffy, B., et al. (2018). Alterations in SCAI Expression during Cell Plasticity, Fibrosis and Cancer. Pathol. Oncol. Res. 24, 641–651.

116. Bhattacharya, C., Wang, X., and Becker, D. (2012). The DEAD/DEAH box helicase, DDX11, is essential for the survival of advanced melanomas. Mol. Cancer 11, 82.

117. Li, J., Liu, L., Liu, X., Xu, P., Hu, Q., and Yu, Y. (2019). The Role of Upregulated DDX11 as A Potential Prognostic and Diagnostic Biomarker in Lung Adenocarcinoma. J. Cancer 10, 4208– 4216.

118. Mahtab, M., Boavida, A., Santos, D., and Pisani, F.M. (2021). The Genome Stability Maintenance DNA Helicase DDX11 and Its Role in Cancer. Genes 12. 10.3390/genes12030395.

119. Marchese, F.P., Grossi, E., Marín-Béjar, O., Bharti, S.K., Raimondi, I., González, J., Martínez-Herrera, D.J., Athie, A., Amadoz, A., Brosh, R.M., Jr, et al. (2016). A Long Noncoding RNA Regulates Sister Chromatid Cohesion. Mol. Cell 63, 397–407.

120. Patton, E.E., Mueller, K.L., Adams, D.J., Anandasabapathy, N., Aplin, A.E., Bertolotto, C., Bosenberg, M., Ceol, C.J., Burd, C.E., Chi, P., et al. (2021). Melanoma models for the next generation of therapies. Cancer Cell 39, 610–631.

121. Rodriguez, H., Zenklusen, J.C., Staudt, L.M., Doroshow, J.H., and Lowy, D.R. (2021). The next horizon in precision oncology: Proteogenomics to inform cancer diagnosis and treatment. Cell 184, 1661–1670.

122. Betancourt, L.H., Sanchez, A., Pla, I., Kuras, M., Zhou, Q., Andersson, R., and Marko-Varga, G. (2018). Quantitative Assessment of Urea In-Solution Lys-C/Trypsin Digestions Reveals Superior Performance at Room Temperature over Traditional Proteolysis at 37 °C. J. Proteome Res. 17, 2556–2561.

123. Murillo, J.R., Kuras, M., Rezeli, M., Miliotis, T., Betancourt, L., and Marko-Varga, G. (2018). Automated phosphopeptide enrichment from minute quantities of frozen malignant melanoma tissue. PLoS One 13, e0208562.

124. Tyanova, S., Temu, T., Sinitcyn, P., Carlson, A., Hein, M.Y., Geiger, T., Mann, M., and Cox, J. (2016). The Perseus computational platform for comprehensive analysis of (prote)omics data. Nat. Methods 13, 731–740.

125. Kuras, M., Betancourt, L.H., Rezeli, M., Rodriguez, J., Szasz, M., Zhou, Q., Miliotis, T., Andersson, R., and Marko-Varga, G. (2019). Assessing Automated Sample Preparation Technologies for High-Throughput Proteomics of Frozen Well Characterized Tissues from Swedish Biobanks. J. Proteome Res. 18, 548–556.

126. Bruderer, R., Bernhardt, O.M., Gandhi, T., Miladinović, S.M., Cheng, L.-Y., Messner, S., Ehrenberger, T., Zanotelli, V., Butscheid, Y., Escher, C., et al. (2015). Extending the limits of quantitative proteome profiling with data-independent acquisition and application to acetaminophen-treated three-dimensional liver microtissues. Mol. Cell. Proteomics 14, 1400– 1410.

127. Liu, W., Payne, S.H., Ma, S., and Fenyö, D. (2019). Extracting Pathway-level Signatures from Proteogenomic Data in Breast Cancer Using Independent Component Analysis. Mol. Cell. Proteomics 18, S169–S182.

128. Linding, R., Jensen, L.J., Pasculescu, A., Olhovsky, M., Colwill, K., Bork, P., Yaffe, M.B., and Pawson, T. (2008). NetworKIN: a resource for exploring cellular phosphorylation networks. Nucleic Acids Res. 36, D695–D699.

129. Miller, M.L., Jensen, L.J., Diella, F., Jørgensen, C., Tinti, M., Li, L., Hsiung, M., Parker, S.A., Bordeaux, J., Sicheritz-Ponten, T., et al. (2008). Linear motif atlas for phosphorylation-dependent signaling. Sci. Signal. 1, ra2.

130. Wei, W., Sun, Z., da Silveira, W.A., Yu, Z., Lawson, A., Hardiman, G., Kelemen, L.E., and Chung, D. (2019). Semi-supervised identification of cancer subgroups using survival outcomes and overlapping grouping information. Stat. Methods Med. Res. 28, 2137–2149.

131. Betancourt, L.H., Szasz, A.M., Kuras, M., Rodriguez Murillo, J., Sugihara, Y., Pla, I., Horvath, Z., Pawłowski, K., Rezeli, M., Miharada, K., et al. (2019). The Hidden Story of Heterogeneous B-raf V600E Mutation Quantitative Protein Expression in Metastatic Melanoma-Association with Clinical Outcome and Tumor Phenotypes. Cancers 11. 10.3390/cancers11121981.

132. Huang, P.-J., Lee, C.-C., Tan, B.C.-M., Yeh, Y.-M., Julie Chu, L., Chen, T.-W., Chang, K.-P., Lee, C.-Y., Gan, R.-C., Liu, H., et al. CMPD. the Cancer Mutant Proteome Database. http://cgbc.cgu.edu.tw/cmpd/.

133. Zhu, Y., Orre, L.M., Johansson, H.J., Huss, M., Boekel, J., Vesterlund, M., Fernandez-Woodbridge, A., Branca, R.M.M., and Lehtiö, J. (2018). Discovery of coding regions in the human genome by integrated proteogenomics analysis workflow. Nat. Commun. 9, 903.

134. Yu, G., Wang, L.-G., Han, Y., and He, Q.-Y. (2012). clusterProfiler: an R package for comparing biological themes among gene clusters. OMICS 16, 284–287.

135. KEGG API Kyoto Encyclopedia of Genes and Genomes. http://rest.kegg.jp/link/hsa/pathway.

136. KEGG Mapper – Convert ID https://www.genome.jp/kegg/tool/conv_id.html.

137. Kanehisa, M., and Goto, S. (2000). KEGG: kyoto encyclopedia of genes and genomes. Nucleic Acids Res. 28, 27–30.

138. Kanehisa, M. (2019). Toward understanding the origin and evolution of cellular organisms. Protein Sci. 28, 1947–1951.

139. Kanehisa, M., Furumichi, M., Sato, Y., Ishiguro-Watanabe, M., and Tanabe, M. (2021). KEGG: integrating viruses and cellular organisms. Nucleic Acids Res. 49, D545–D551.

140. Desiere, F., Deutsch, E.W., King, N.L., Nesvizhskii, A.I., Mallick, P., Eng, J., Chen, S., Eddes, J., Loevenich, S.N., and Aebersold, R. (2006). The PeptideAtlas project. Nucleic Acids Res. 34, D655–D658.

141. Li, J., Duncan, D.T., and Zhang, B. (2010). CanProVar: a human cancer proteome variation database. Hum. Mutat. 31, 219–228.

142. Zhang, M., Wang, B., Xu, J., Wang, X., Xie, L., Zhang, B., Li, Y., and Li, J. (2017). CanProVar 2.0: An Updated Database of Human Cancer Proteome Variation. J. Proteome Res. 16, 421–432.

143. Durinck, S., Moreau, Y., Kasprzyk, A., Davis, S., De Moor, B., Brazma, A., and Huber, W. (2005). BioMart and Bioconductor: a powerful link between biological databases and microarray data analysis. Bioinformatics 21, 3439–3440.

144. Durinck, S., Spellman, P.T., Birney, E., and Huber, W. (2009). Mapping identifiers for the integration of genomic datasets with the R/Bioconductor package biomaRt. Nat. Protoc. 4, 1184– 1191.

145. Sondka, Z., Bamford, S., Cole, C.G., Ward, S.A., Dunham, I., and Forbes, S.A. (2018). The COSMIC Cancer Gene Census: describing genetic dysfunction across all human cancers. Nat. Rev. Cancer 18, 696–705.

146. Cosmic (2022). COSMIC. 10.1093/nar/gkw1121.

147. Phan, L., Jin, Y., Zhang, H., Qiang, W., Shekhtman, E., Shao, D., Revoe, D., Villamarin, R., Ivanchenko, E., Kimura, M., et al. (2020). ALFA: Allele Frequency Aggregator.

148. Lek, M., Karczewski, K.J., Minikel, E.V., Samocha, K.E., Banks, E., Fennell, T., O’Donnell-Luria, A.H., Ware, J.S., Hill, A.J., Cummings, B.B., et al. (2016). Analysis of protein-coding genetic variation in 60,706 humans. Nature 536, 285–291.

149. 1000 Genomes Project Consortium, Auton, A., Brooks, L.D., Durbin, R.M., Garrison, E.P., Kang, H.M., Korbel, J.O., Marchini, J.L., McCarthy, S., McVean, G.A., et al. (2015). A global reference for human genetic variation. Nature 526, 68–74.

150. International HapMap Consortium (2003). The International HapMap Project. Nature 426, 789–796.

151. Blumenberg, L., Kawaler, E.A., Cornwell, M., Smith, S., Ruggles, K.V., and Fenyö, D. (2021). BlackSheep: A Bioconductor and Bioconda Package for Differential Extreme Value Analysis. J. Proteome Res. 20, 3767–3773.

152. Yuan, Y., Van Allen, E.M., Omberg, L., Wagle, N., Amin-Mansour, A., Sokolov, A., Byers, L.A., Xu, Y., Hess, K.R., Diao, L., et al. (2014). Assessing the clinical utility of cancer genomic and proteomic data across tumor types. Nat. Biotechnol. 32, 644–652.

153. Ogata, Y., Heppelmann, C.J., Heppelmann, C.J., Charlesworth, M.C., Madden, B.J., Miller, M.N., Kalli, K.R., Cliby, W.A., Bergen, H.R., 3rd, Saggese, D.A., et al. (2006). Elevated levels of phosphorylated fibrinogen-alpha-isoforms and differential expression of other post-translationally modified proteins in the plasma of ovarian cancer patients. J. Proteome Res. 5, 3318–3325.

154. Nagel, T., Klaus, F., Ibanez, I.G., Wege, H., Lohse, A., and Meyer, B. (2018). Fast and facile analysis of glycosylation and phosphorylation of fibrinogen from human plasma-correlation with liver cancer and liver cirrhosis. Anal. Bioanal. Chem. 410, 7965–7977.

155. Guida, M., Ravaioli, A., Sileni, V.C., Romanini, A., Labianca, R., Freschi, A., Brugnara, S., Casamassima, A., Lorusso, V., Nanni, O., et al. (2003). Fibrinogen: a novel predictor of responsiveness in metastatic melanoma patients treated with bio-chemotherapy: IMI (italian melanoma inter-group) trial. J. Transl. Med. 1, 13.

156. Ciereszko, A., Dietrich, M.A., Słowińska, M., Nynca, J., Ciborowski, M., Kisluk, J., Michalska- Falkowska, A., Reszec, J., Sierko, E., and Nikliński, J. (2019). Identification of protein changes in the blood plasma of lung cancer patients subjected to chemotherapy using a 2D-DIGE approach. PLoS One 14, e0223840.

157. Gunji, Y., and Gorelik, E. (1988). Role of fibrin coagulation in protection of murine tumor cells from destruction by cytotoxic cells. Cancer Res. 48, 5216–5221.

158. Cardinali, M., Uchino, R., and Chung, S.I. (1990). Interaction of fibrinogen with murine melanoma cells: covalent association with cell membranes and protection against recognition by lymphokine-activated killer cells. Cancer Res. 50, 8010–8016.

159. Palumbo, J.S., Kombrinck, K.W., Drew, A.F., Grimes, T.S., Kiser, J.H., Degen, J.L., and Bugge, T.H. (2000). Fibrinogen is an important determinant of the metastatic potential of circulating tumor cells. Blood 96, 3302–3309.

160. Ryu, J.K., Petersen, M.A., Murray, S.G., Baeten, K.M., Meyer-Franke, A., Chan, J.P., Vagena, E., Bedard, C., Machado, M.R., Rios Coronado, P.E., et al. (2015). Blood coagulation protein fibrinogen promotes autoimmunity and demyelination via chemokine release and antigen presentation. Nat. Commun. 6, 8164.

161. Furfaro, A.L., Ottonello, S., Loi, G., Cossu, I., Piras, S., Spagnolo, F., Queirolo, P., Marinari, U.M., Moretta, L., Pronzato, M.A., et al. (2020). HO-1 downregulation favors BRAFV600 melanoma cell death induced by Vemurafenib/PLX4032 and increases NK recognition. Int. J. Cancer 146, 1950–1962.

162. Was, H., Cichon, T., Smolarczyk, R., Rudnicka, D., Stopa, M., Chevalier, C., Leger, J.J., Lackowska, B., Grochot, A., Bojkowska, K., et al. (2006). Overexpression of heme oxygenase-1 in murine melanoma: increased proliferation and viability of tumor cells, decreased survival of mice. Am. J. Pathol. 169, 2181–2198.

163. Hjortsø, M.D., and Andersen, M.H. (2014). The expression, function and targeting of haem oxygenase-1 in cancer. Curr. Cancer Drug Targets 14, 337–347.

164. Jasmer, K.J. (2015). The role of heme oxygenase in metastatic melanoma tumorigenicity.

165. Yanagisawa, M., Huveldt, D., Kreinest, P., Lohse, C.M., Cheville, J.C., Parker, A.S., Copland, J.A., and Anastasiadis, P.Z. (2008). A p120 catenin isoform switch affects Rho activity, induces tumor cell invasion, and predicts metastatic disease. J. Biol. Chem. 283, 18344–18354.

166. Kourtidis, A., Yanagisawa, M., Huveldt, D., Copland, J.A., and Anastasiadis, P.Z. (2015). Pro-Tumorigenic Phosphorylation of p120 Catenin in Renal and Breast Cancer. PLoS One 10, e0129964.

167. Aho, S., Levänsuo, L., Montonen, O., Kari, C., Rodeck, U., and Uitto, J. (2002). Specific sequences in p120ctn determine subcellular distribution of its multiple isoforms involved in cellular adhesion of normal and malignant epithelial cells. J. Cell Sci. 115, 1391–1402.

168. van Hengel, J., and van Roy, F. (2007). Diverse functions of p120ctn in tumors. Biochim. Biophys. Acta 1773, 78–88.

169. Aslund-Ostberg, A.M., Marklund, B., and Hegen, C. (1992). [Outpatient clinics, open during evening hours for consultation on skin changes, attracted many visitors]. Lakartidningen 89, 3923–3924.

170. Pieters, T., van Roy, F., and van Hengel, J. (2012). Functions of p120ctn isoforms in cell-cell adhesion and intracellular signaling. Front. Biosci. 17, 1669–1694.

171. Ren, S., Chai, L., Wang, C., Li, C., Ren, Q., Yang, L., Wang, F., Qiao, Z., Li, W., He, M., et al. (2015). Human malignant melanoma-derived progestagen-associated endometrial protein immunosuppresses T lymphocytes in vitro. PLoS One 10, e0119038.

172. Liu, S. (2011). Editorial - a potential new target gene of the master-regulator microphthalmia-associated transcription factor in melanoma. Ochsner J. 11, 210–211.

173. Ren, S., Howell, P.M., Jr, Han, Y., Wang, J., Liu, M., Wang, Y., Quan, G., Du, W., Fang, L., and Riker, A.I. (2011). Overexpression of the progestagen-associated endometrial protein gene is associated with microphthalmia-associated transcription factor in human melanoma. Ochsner J. 11, 212–219.

174. Ren, S., Liu, S., Howell, P.M., Jr, Zhang, G., Pannell, L., Samant, R., Shevde-Samant, L., Tucker, J.A., Fodstad, O., and Riker, A.I. (2010). Functional characterization of the progestagen- associated endometrial protein gene in human melanoma. J. Cell. Mol. Med. 14, 1432–1442.

175. Luke, J.J., Flaherty, K.T., Ribas, A., and Long, G.V. (2017). Targeted agents and immunotherapies: optimizing outcomes in melanoma. Nat. Rev. Clin. Oncol. 14, 463–482.

176. Gao, X., Leone, G.W., and Wang, H. (2020). Cyclin D-CDK4/6 functions in cancer. Adv. Cancer Res. 148, 147–169.

177. Kwong, L.N., and Davies, M.A. (2013). Navigating the therapeutic complexity of PI3K pathway inhibition in melanoma. Clin. Cancer Res. 19, 5310–5319.

178. Yuan, T.L., and Cantley, L.C. (2008). PI3K pathway alterations in cancer: variations on a theme. Oncogene 27, 5497–5510.

179. Lord, C.C., Thomas, G., and Brown, J.M. (2013). Mammalian alpha beta hydrolase domain (ABHD) proteins: Lipid metabolizing enzymes at the interface of cell signaling and energy metabolism. Biochim. Biophys. Acta 1831, 792–802.

180. Huang, L., Chen, J., Zhao, Y., Gu, L., Shao, X., Li, J., Xu, Y., Liu, Z., and Xu, Q. (2019). Key candidate genes of STAT1 and CXCL10 in melanoma identified by integrated bioinformatical analysis. IUBMB Life 71, 1634–1644.

