## Supplementary Figures S1-S5 for "Histopathology-assisted proteogenomics provides foundations for stratification of melanoma metastases"

### SUPPLEMENTARY FIGURES AND TABLES

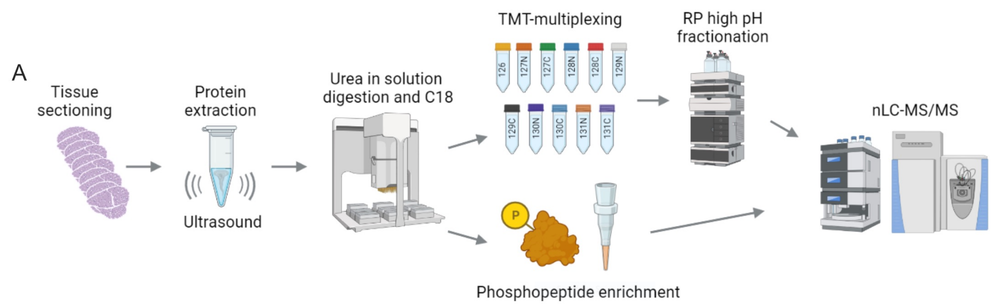


B

| Age at diagnosis | | Survival from sample collection | | Overall survival | |
|---|---|---|---|---|---|
| Range | 23–96 years | Range | 0.1–25.2 years | Range | 0.2–42.5 years |
| Median ± SD | 64 ± 13.9 years | Median ± SD | 2 ± 5.1 years | Median ± SD | 4.9 ± 7 years |
| **Gender** | | **Disease stage (patients)** | | **Localization** | |
| Female | 47 | I-II | 3 | Lymph node | 121 |
| Male | 87 | III | 97 | Cutaneous | 1 |
| NA | 3 | IV | 34 | Subcutaneous | 7 |
| | | NA | 3 | Visceral | 3 |
| | | | | NA | 5 |
| **Mutational status** | | **Type of BRAF mutation** | | **Status/Remarks** | |
| wt | 36 | V600E | 45 | Alive | 26 |
| BRAF | 49 | V600K | 2 | Dead | 91 |
| NRAS | 35 | V600R | 1 | Unknown cause | 16 |
| NA | 17 | V600A | 1 | NA | 4 |

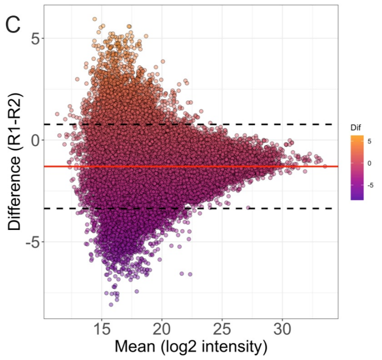


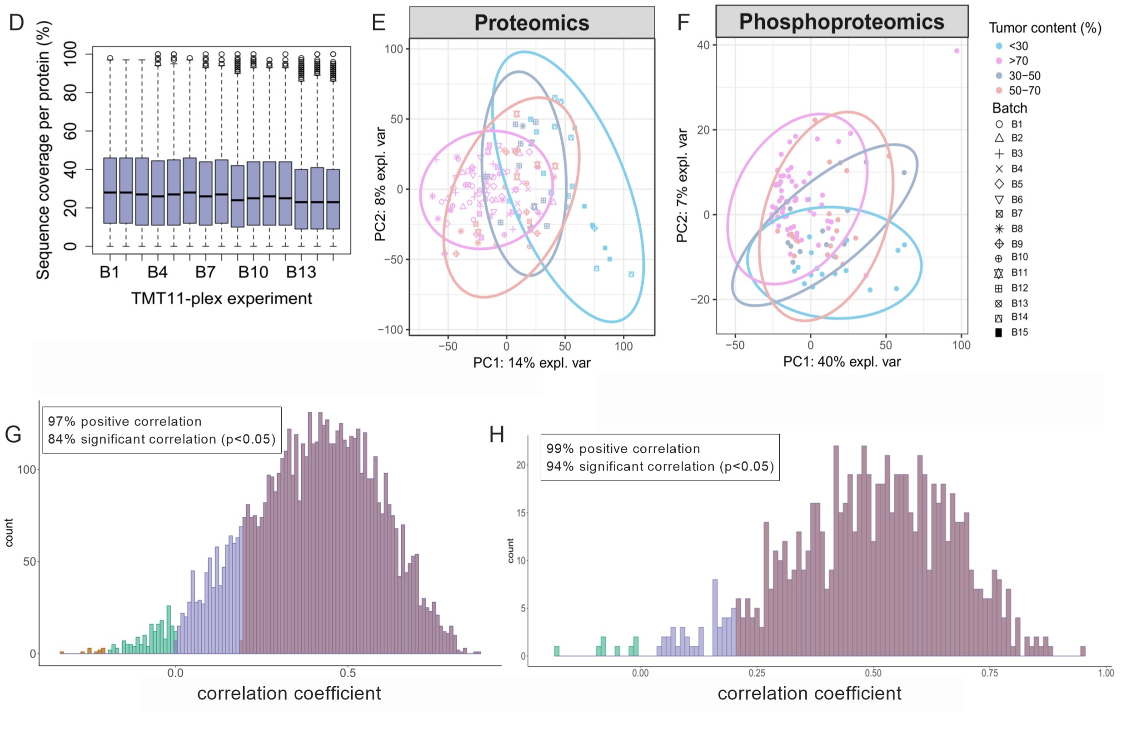


**Figure S1. Proteogenomics of melanoma metastases: workflow, cohort and quality control of the omics data**
(A) Global Proteomic and phosphoproteomic workflow.
(B) Summary table of clinical features in the discovery cohort.
(C) Bland-Altman plot depicting the agreement across repeated experiments of batch one (B1), upper limit: 0.770, lower limit: -3.363, and mean difference: -1.297.

(D) Distribution of sequence coverage of the identified proteins by MS/MS across the fifteen TMT11 plex batches (whiskers show the 5–95 percentiles, and the dots represent the outliers).
(E) Principal component analysis of the TMT global proteome data after normalization and ratio calculation. The ellipses represent the 95% confidence interval per group based on 8,125 proteins.
(F) Principal component analysis of 1,267 commonly quantified phosphosites. The ellipses represent the 95% confidence interval per group.
(G) mRNA and protein abundance correlation (median 0.408), where 84% of the mRNA and protein pairs (6,101) showed a significant correlation (p-value 0.05) across 104 patient samples.
(H) Protein and phosphoprotein abundance correlation (median 0.506), where 94% of the proteins and phosphoprotein pairs (809) showed a significant correlation (p-value 0.05) across 94 patient samples.

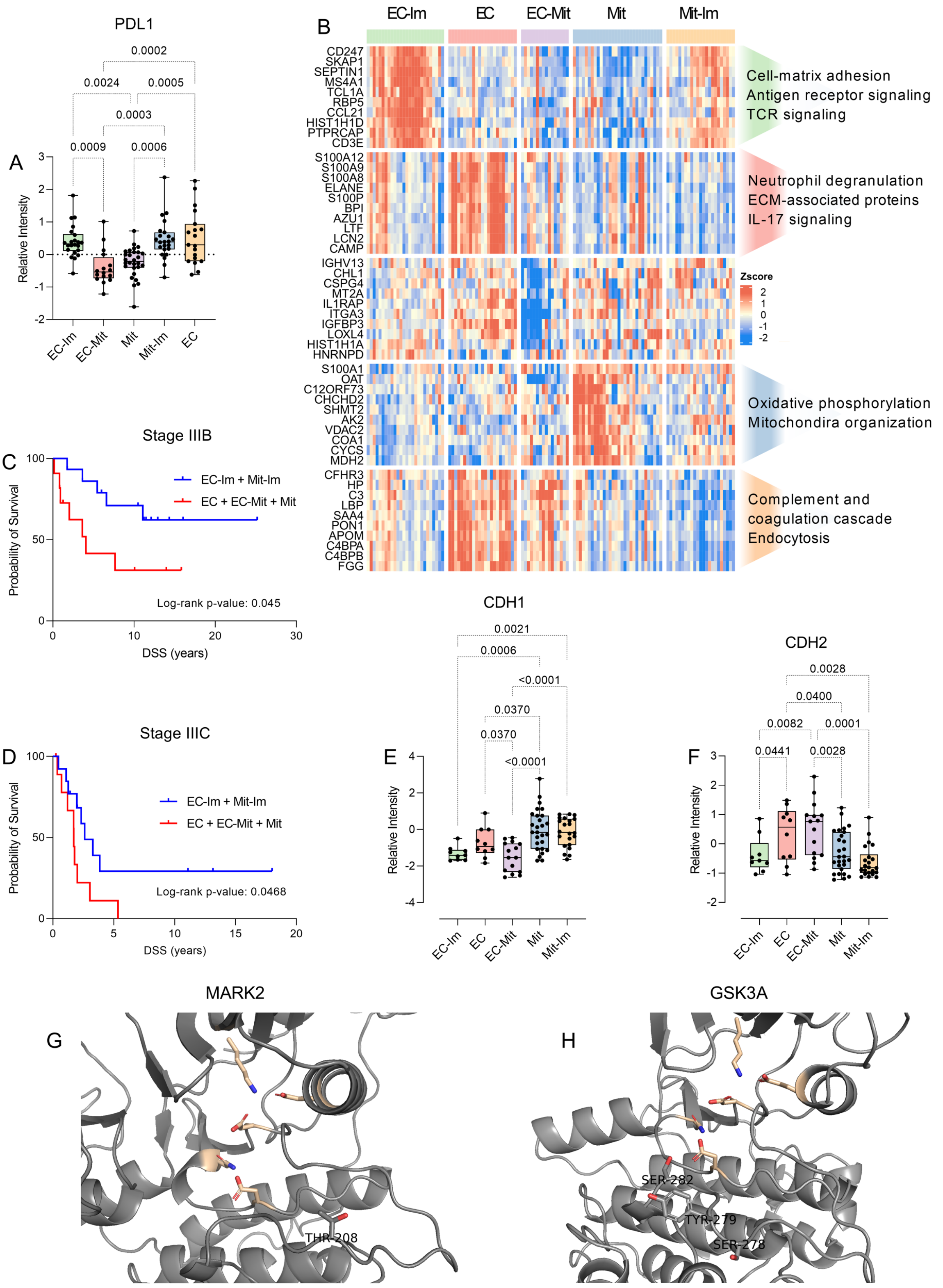


**Figure S2. Molecular and clinical features of the proteomic subtypes**
(A) PDL1 protein expression across the five proteomic subtypes.
(B) The ten proteins most strongly contributing to the ICs correlated with the proteomic subtypes and their enriched pathways (FDR < 0.05).

(C) Disease-specific survival probability for patients in disease stage IIIB, divided into subtypes associated with long and short survival.
(D) Disease-specific survival probability for patients in disease stage IIIC, divided into subtypes associated with long and short survival.
(E) Protein expression of CDH1 across the five proteomic subtypes.
(F) Protein expression of CDH2 across the five proteomic subtypes.
(G) Location of regulated phosphosites in the activation loops within three-dimensional structures of the MARK2 kinase.
(H) Location of regulated phosphosites in the activation loops within three-dimensional structures of the GSK3A kinases.

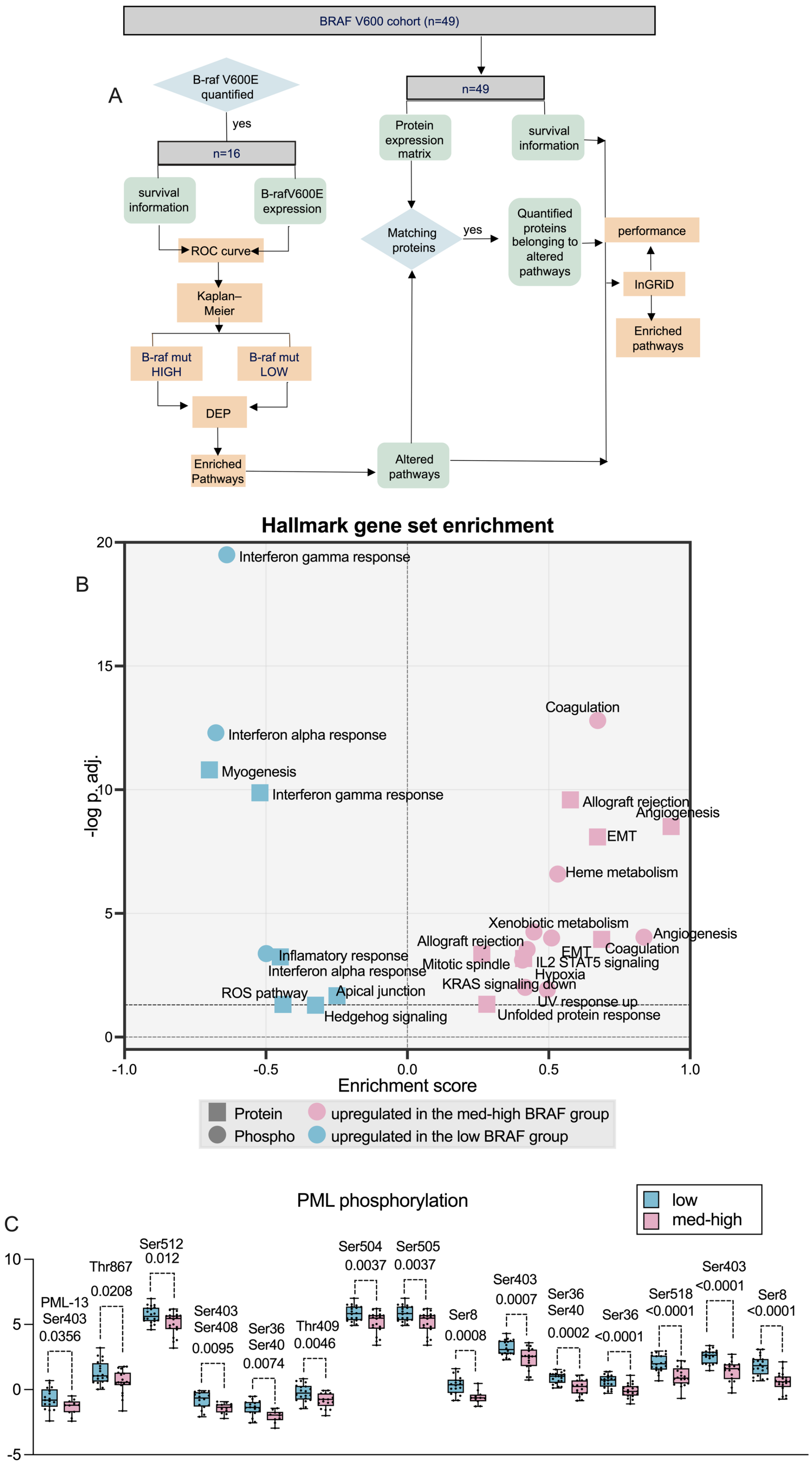
A
BRAF V600 cohort (n=49)
B-raf V600E quantified
yes
n=16
survival information
B-rafV600E expression
ROC curve
Kaplan–Meier
B-raf mut HIGH
B-raf mut LOW
DEP
Enriched Pathways
n=49
Protein expression matrix
survival information
Matching proteins
yes
Quantified proteins belonging to altered pathways
performance
InGRiD
Enriched pathways
Altered pathways
B
Hallmark gene set enrichment
-log p.adj.
Enrichment score
Interferon gamma response
Coagulation
Interferon alpha response
Myogenesis
Interferon gamma response
Allograft rejection
Angiogenesis
EMT
Heme metabolism
Xenobiotic metabolism
Angiogenesis
Allograft rejection
EMT
Coagulation
Inflamatory response
Mitotic spindle
IL2 STAT5 signaling
Interferon alpha response
Hypoxia
KRAS signaling down
ROS pathway
Apical junction
UV response up
Hedgehog signaling
Unfolded protein response
Protein
Phospho
upregulated in the med-high BRAF group
upregulated in the low BRAF group
C
PML phosphorylation
low
med-high
PML-13 Ser403 0.0356
Thr867 0.0208
Ser512 0.012
Ser403 Ser408 0.0095
Ser36 Ser40 0.0074
Thr409 0.0046
Ser504 0.0037
Ser505 0.0037
Ser8 0.0008
Ser403 0.0007
Ser36 Ser40 0.0002
Ser36 <0.0001
Ser518 <0.0001
Ser403 <0.0001
Ser8 <0.0001

**Figure S3. Molecular difference between the low and med-high BRAF V600 mortality risk groups**

(A) Workflow for the association between BRAF V600E expression and patient survival in a larger cohort. The orange squares indicate statistical tests.

(B)Enrichment of the differentially expressed proteins (p-value 0.05) and phosphosites between the low and medium-high mortality risk groups using the Hallmark gene set.

(C) Phosphosite expression of the PML protein and the PML-13 isoform, between the low and medium-high mortality risk groups.

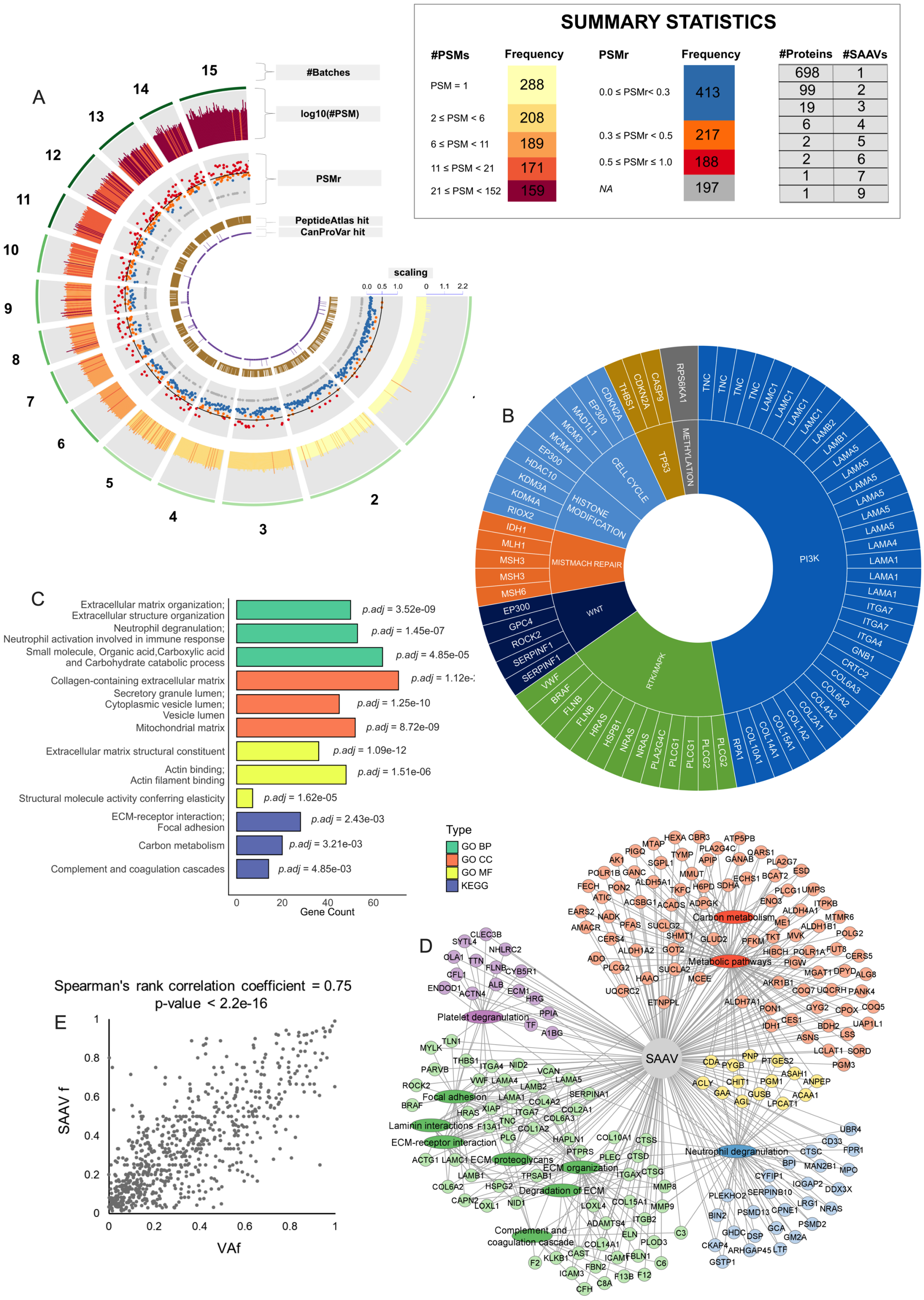

**Figure S4. Overview of identified SAAVs**

(A) Summary of the 1,015 validated SAAVs, including the number of TMT batches, number of PSMs, SAAVf, and the availability in PeptideAtlas and CanProVar databases. The embedded summary statistics table includes bar charts showing the number of PSMs (#PSMs), the SAAVf, and the distribution of the number of SAAVs per protein.

(B) Identified Genes with SAAVs belonging to signaling pathways frequently dysregulated in melanoma.

(C) KEGG and GO enrichment analysis of the 828 proteins with SAAVs.

(D) Network of proteins with SAAVs linked to KEGG and GO-enriched pathways

(E) Significant correlation between of SAAVf and VAf

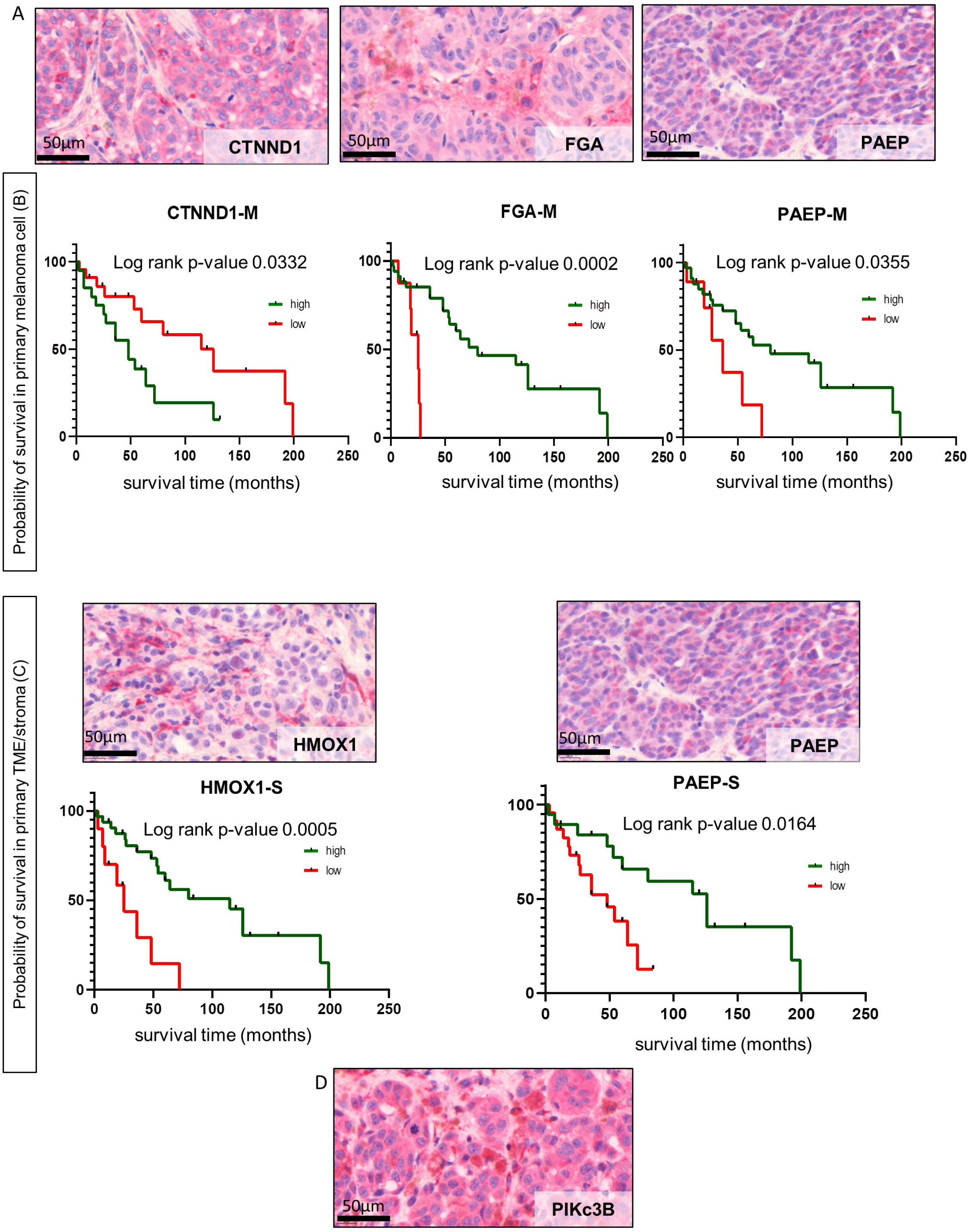


**Figure S5. Different expressional features of the candidate biomarkers and survival during progression**

A) Tissue heterogeneity of CTNND1, FGA, and PAEP protein expression in melanoma tissue. IHC, fast red colorimetry – OM 112x; scale bar 50µm.

B) Kaplan-Meier analyses of the OS rates for patients in association with high (green) and low (red) expression of the markers (CTNND1, PAEP, FGA) in melanoma cells with significant differences.

C) IHC expression and Kaplan-Meier analyses of the OS rates for patients with high (green) and low (red) expression of HMOX1 and PAEP in the TME with significant differences. IHC, fast red colorimetry – OM 112x; scale bar 50µm.
D) Tissue heterogeneity of PIKc3B protein IHC expression.

**Tables**

**Table S1. Clinical and histopathology data of melanoma cohorts**
**Table S2. Proteogenomic data**
**Table S3. Top enriched GO and KEGG in 3000 proteins for subtypes and Clinico-histopathological feature associations**
**Table S4. OMICs and Independent Component Analysis**
**Table S5. BRAF survival risk analysis**
**Table S6. Single Amino Acid Variant analysis**
**Table S7. Protein and mRNA profiles in the tumor microenvironment (TME)**
**Table S8. Survival analysis**
**Table S9. Immunohistochemistry analysis of survival markers**
